# Liebenberg syndrome severity arises from variations in *Pitx1* locus topology and ectopically transcribing cells

**DOI:** 10.1101/2024.03.04.582675

**Authors:** Olimpia Bompadre, Raquel Rouco, Fabrice Darbellay, Antonella Rauseo, Fanny Guerard-Millet, Claudia Gentile, Marie Kmita, Guillaume Andrey

## Abstract

Enhancer hijacking, a common cause of gene misregulation linked to disease, occurs when non-matching enhancers and promoters interact ectopically. This interaction is made possible by genetic changes that alter the arrangement or insulation of gene regulatory landscapes. While the concept of enhancer hijacking is well understood, the specific reasons behind the variation in phenotypic severity or the point at which those phenotypes become evident remain unexplored. In this work, we expand on the ectopic activation of the hindlimb-specific transcription factor *Pitx1* by one of its own enhancers, *Pen*, in forelimb tissues that causes the Liebenberg syndrome. We combine a previously developed *in-embryo* cell-tracing approach to a series of inversions and relocations to show that reduction in *Pitx1*-*Pen* relative genomic positioning leads to increased proportions of *Pitx1* forelimb-expressing cells and more severe phenotypical outcomes. We demonstrate that the *Pitx1* locus assumes an active topology when enhancer-promoter contacts are required for transcription and that its promoter generates consistent transcription levels across different alleles. Finally, we show that changes in 3D chromatin structure and enhancer-promoter contacts are not the result of *Pitx1* transcriptional activity. In summary, our work shows that variation in enhancer-promoter interactions can lead to pathogenic locus activation in variable proportions of cells which, in turn, define phenotypic severity.

## Introduction

The restriction of enhancer-promoter contacts is a fundamental feature of gene regulation. This was shown to be mediated by domains of preferential interactions called topologically-associating domains (TADs). Indeed, TADs foster high internal chromatin interactions while reducing interactions with external regions. Biophysically, TADs are believed to be formed by a process called loop extrusion where cohesin molecules extrude chromatin until reaching CTCF which induces a temporary stalling of the process (Fudenberg et al., 2016; Sanborn et al., 2015). Changes in CTCF binding therefore impact the 3D architecture of loci and enhancer-promoter contacts (Despang et al., 2019). Moreover, tissue-specific chromatin interactions can actively control enhancer-promoter communications in a spatiotemporally-defined manner, enabling the activation of associated genes (Andrey et al., 2013; Deng et al., 2012; Deng et al., 2014; Kragesteen et al., 2018).

Alterations in this organized process can lead to the wrongful connection between non-matching enhancers and promoters, leading to gene de-repression and expression in ectopic tissues, in a process named “enhancer-highjacking”. In particular, structural variants (SVs) that impact the topological organisation of loci have been shown to lead to congenital malformations in such a way (Franke et al., 2016; Lupianez et al., 2015; Spielmann et al., 2018). Although the *patho*-mechanism of SV-induced enhancer-hijacking has been documented across numerous loci, these accounts often overlook the influence of variations in SV breakpoints on disease outcomes or severity (Zaugg et al., 2022). Furthermore, the precise relationship between distinct SVs and subsequent changes in the 3D genome architecture, chromatin modifications, and ectopic gene transcription is yet to be fully elucidated.

This is what happens at the *Pitx1* locus, where different SVs underlying the Liebenberg syndrome, a congenital malformation associated to a partial arm-to-leg transformation, are associated with variable morphological changes (Al-Qattan et al., 2013; Kragesteen et al., 2019; Seoighe et al., 2014; Spielmann et al., 2012). During normal development, the *Pitx1* gene is specifically expressed in developing hindlimb, and not in forelimbs, where it controls hindlimb outgrowth and differentiation into a leg (Infante et al., 2013; Lanctot et al., 1997; Nemec et al., 2017). So far, three limb enhancers have been identified at the locus: *PelB, RA4* and *Pen* (Kragesteen et al., 2018; Thompson et al., 2018). Notably, another enhancer, *PDE,* has been described to contact the gene and as being strongly marked with H3K27ac in hindlimb, however, in reporter assays, the region only displays activity in the developing mandible (Kragesteen et al., 2018; Sarro et al., 2018). Importantly, both *RA4* and *Pen* display a fore- and hindlimb activity when assayed in transgenic reporter approaches, and indeed, in the Liebenberg syndrome, the *Pitx1* gene gets *endo*-activated, i.e. ectopically activated by one of these two enhancers, *Pen*, in developing forelimbs (Kragesteen et al., 2018). This activation results from SVs that re-arrange the locus and generally bring *Pen*, normally located 400kb away from *Pitx1*, in a closer genetic proximity to *Pitx1*. Patients with SVs that slightly reduce the *Pitx1-Pen* genetic distance show rather mild malformation features, yet, patients where *Pitx1-Pen* linear distance is strongly reduced display more severe ones (**Supplementary Fig. S1**, **Supplementary Table S1**)(Al-Qattan et al., 2013; Kragesteen et al., 2019; Seoighe et al., 2014; Spielmann et al., 2012).

Here, we combine a previously developed *in-embryo* cell-tracing approach with engineered Liebenberg structural variants and *Pen* relocations to measure and isolate *Pitx1*-expressing cells in mouse forelimbs (Rouco et al., 2021). In this context, we explore how structural variants can cause different degrees of phenotypic manifestations by identifying their link to ectopically expressing cells and transcriptional activities. Moreover, we investigate how de-repression or targeted activation of *Pitx1* can impact transcriptional activities and the locus topology.

## Results

### *Pitx1-Pen* relative genomic position affects the proportion of *Pitx1* ectopically expressing cells

To address how differential SVs breakpoints lead to gene mis-activation, we took advantage of the previously described *Pitx1^EGFP^* sensor allele that allows for the tracking and sorting of *Pitx1* active and inactive cells from developing tissues (Rouco et al, 2021). We re-engineered in the *Pitx1^EGFP^* background a previously published inversion leading to Liebenberg syndrome in mice: *Pitx1^EGFP;Inv1+/-^*, as well a larger one *Pitx1^EGFP;Inv2+/-^* (**Fig. 1A, B, C**) (Kragesteen et al., 2018). These inversions place *Pen* at the positions of *RA4* and *PDE*, located 225kb and 116 kb from *Pitx1*, respectively. Using Capture-HiC (C-HiC) in *Pitx1^EGFP^*mouse Embryonic Stem Cells (mESCs) we could observe at both integration sites higher contact frequencies with *Pitx1*, than at *Pen*, with 1.1x at *RA4* and 2.7x at *PDE* (**Fig. 1A**). To measure how inversions perturb the locus poised 3D organisation, we performed C-HiC in *Pitx1^EGFP;Inv1+/-^*and *Pitx1^EGFP;Inv2+/-^* mESCs. In *Pitx1^EGFP;Inv1+/-^*, we observed a similar structure as in control mESCs. In contrast, in *Pitx1^EGFP;Inv2+/-^*, we observed several differences in the locus topology, with increased contact between *Pitx1*, *Pen*, and *Neurog1* (**Fig. 1C**).

**Figure 1:**
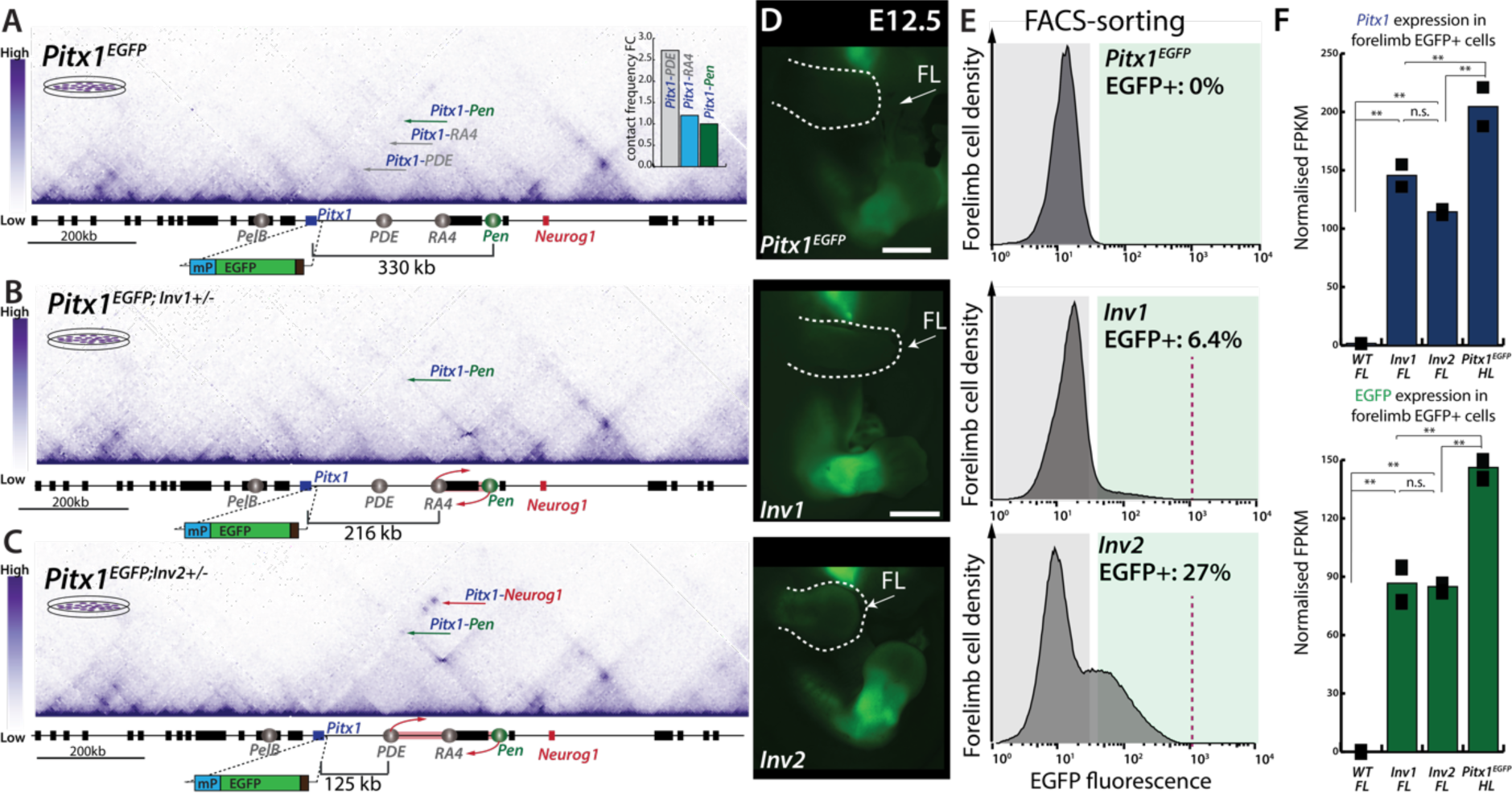
Inversions at the Pitx1 locus lead to increased mis-activation of the gene. **A.** C-HiC analysis of the Pitx1 locus in Pitx1^EGFP^ mESCs. Upper right corner: quantification of interactions between Pitx1 and RA4/PDE/Pen. **B.** A 113kb inversion (Pitx1^EGFP;Inv1+/-^ (Inv1) that swaps the relative position of Pen and RA4 shows a relative decrease in Pitx1-Pen interactions. **C.** A 204kb inversion Pitx1^EGFP;Inv2+/-^ (Inv2) shows an overall increase of contacts between Pitx1 and Pen **D.** Fluorescence microscopy reveals the mis-expression of EGFP and thus Pitx1 in developing forelimbs of Pitx1^EGFP^ andInv1 and Inv2 E12.5 embryos. Forelimbs (FL) are delineated with a dotted white line and a white arrow. **E.** Histogram of EGFP signal and quantification of proportion of mis-expressing cells. The grey and green areas show the delimitation of gating for EGFP- and EGFP+ cells, respectively, in the three alleles. The dotted red line in histograms indicates the upper limit of fluorescence. **F.** Normalised FPKMs of EGFP and Pitx1 in E12.5 wildtype bulk forelimbs, EGFP+ cells of Inv1 and Inv2 forelimbs and EGFP+ cells of Pitx1^EGFP^ hindlimbs. Note a plateau in Pitx1 and EGFP expression in both inversions, which is significantly lower than in EGFP+ cells from Pitx1^EGFP^ hindlimbs. Adjusted p-values are computed using the Wald-test and Benjamini-Hochberg multiple test correction as implemented by the Deseq2 tool where n.s. is a non-significant difference, *= padj < 0.01, **=padj < 0.001 (n=2) (**Supplementary Table S2**).

We then derived *Pitx1^EGFP^*, *Pitx1^EGFP;Inv1+/-^*and *Pitx1^EGFP;Inv2+/-^* E12.5 embryos through tetraploid complementation and characterised forelimb EGFP fluorescence through microscopy and fluorescence activated cell sorting (FACS) (**Fig. 1D, E**)(Artus and Hadjantonakis, 2011). We could measure in *Pitx1^EGFP;Inv1+/-^* forelimbs 6.4% of EGFP-expressing cells in contrast to 0% in *Pitx1^EGFP^* control (**Fig. 1E**). This number rose to 27% in *Pitx1^EGFP;Inv2+/-^* (**Fig. 1E**), this result suggesting that the variation in SV size can alter the proportion of cells ectopically expressing *Pitx1* in the forelimb.

Interestingly, we observed an upper limit of EGFP fluorescence in both inversions, suggesting that the abundance of EGFP in active cells was similar between alleles (**Fig. 1E**). To confirm that hypothesis, we measured transcription in EGFP+ cells of both *Pitx1^EGFP;Inv1+/-^ Pitx1^EGFP;Inv2+/-^* forelimbs using RNA-seq. We observed similar transcription levels of *Pitx1* and EGFP in both alleles (**Fig. 1F, Supplementary Table S2**). In fact, the ectopic transcriptional activity was only 1.5x lower than the one found in wildtype *Pitx1^EGFP^* EGFP+ cells from hindlimbs (**Fig. 1F**). This weak difference might be the result of the heterozygous state of both inversions in *Pitx1^EGFP^* that would indicate a similar transcriptional activity per allele in forelimb EGFP+ cells compared to wildtype hindlimb EGFP+ cells. Yet, as the inverted intervals of both *Pitx1^EGFP;Inv1+/-^* and *Pitx1^EGFP;Inv2+/-^* contain CTCF sites (**Supplementary Fig. S2**) and other *Pitx1* enhancers, the interpretation of the results can be confounding. Therefore, alternative approaches to solely measure the effect of the relocation of *Pen* were further developed.

### A series of *Pen* relocations induce varying proportions of *Pitx1-*expressing cells

To rule out the positional effect induced by the inverted genomic interval, we devised a parallel approach where we solely re-mobilized the *Pen* enhancer itself in a *Pitx1^EGFP;ΔPen^* homozygous deleted background. Here, we inserted *Pen* at the same locations as in the inversions, at *RA4* (*Pitx1^EGFP;ΔPen^*^;*Rel1+/-*^) and at *PDE*(*Pitx1^EGFP;ΔPen^*^;*Rel2+/-*^)(Fig. 2A, B). Moreover, we also introduced *Pen* 7.7kb upstream of the *Pitx1* promoter (*Pitx1^EGFP;ΔPen^*^;*Rel3+/-*^), in a similar genetic distance (10.5kb enhancer-promoter distance) as the one found in the most severe case of Liebenberg syndrome described (Fig. 2C, **Supplementary Fig. S1)**(Seoighe et al., 2014). Of note, with each relocation reducing the genetic distance between *Pitx1* and *Pen*, there is also a consequent reduction in the number of CTCF binding sites separating these two elements (**Supplementary Fig. S2**).

**Figure 2:**
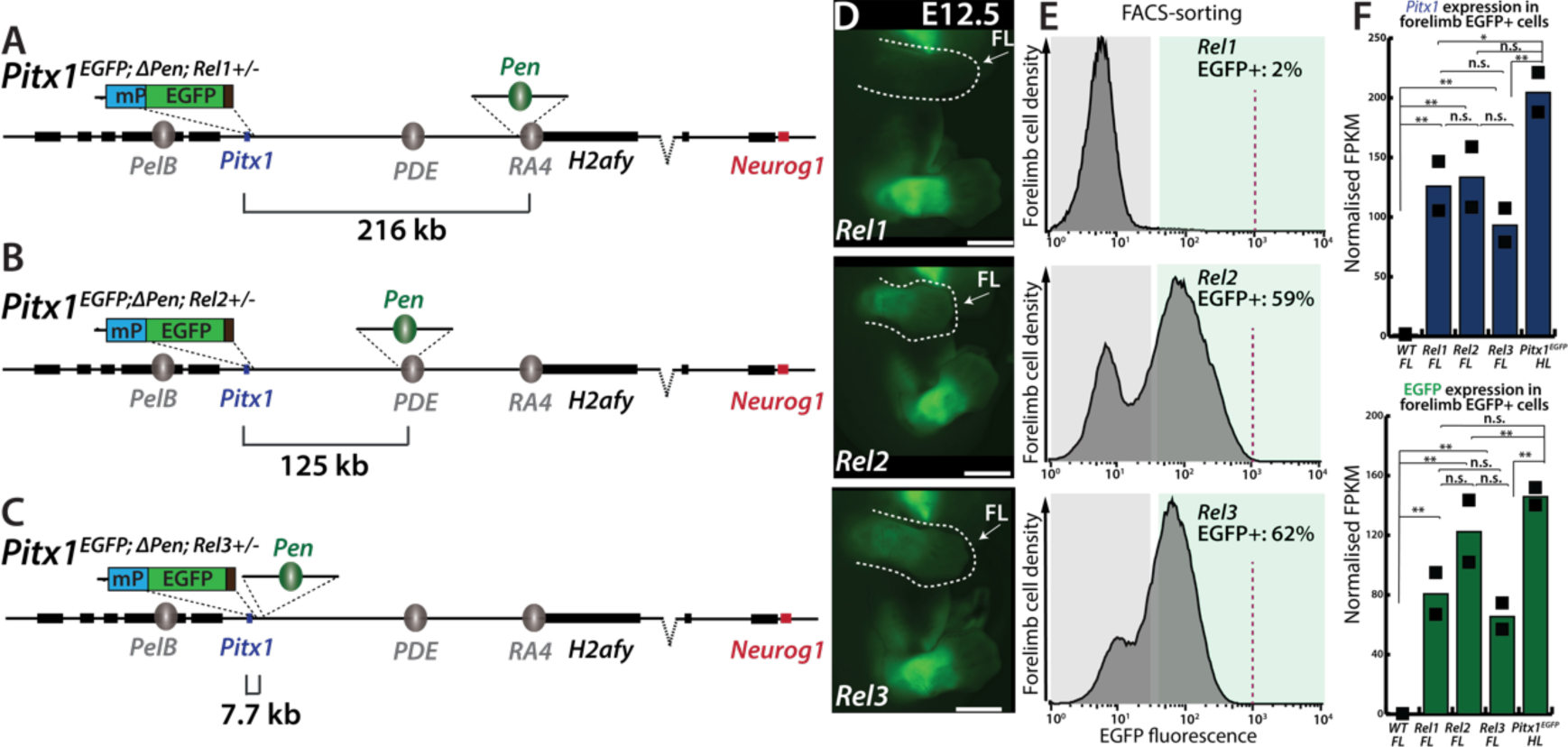
Relocation of Pen through the locus ectopically activates Pitx1. **A**. Illustration of Pitx1^GFP;ΔPen;Rel1+/-^ (Rel1) where Pen is inserted, at RA4, 216kb away to Pitx1. **B**. Illustration of Pitx1^GFP;ΔPen;Rel2+/-^ (Rel2) where Pen is inserted at PDE, 125kb away from Pitx1. **C**. Illustration of Pitx1^GFP;ΔPen;Rel3+/-^ (Rel3) where Pen is inserted 7.7kb away from Pitx1. **D**. Conventional microscopy reveals the mis-expression of EGFP and thus Pitx1 in developing forelimbs of Rel1, Rel2 and Rel3 E12.5 embryos. Forelimbs (FL) are delineated with a dotted white line and a white arrow. **E** Histogram of EGFP signal and quantification of proportion of mis-expressing cells. The grey and green areas show the delimitation of gating for EGFP- and EGFP+ cells, respectively, in the three alleles. The dotted red line in histograms indicates the upper limit of fluorescence. **F**. Normalised FPKMs of EGFP and Pitx1 in E12.5 wildtype bulk forelimbs, EGFP+ cells of Rel1, Rel2 and Rel3 forelimbs and EGFP+ cells of Pitx1^EGFP^ hindlimbs. Note the consistent Pitx1 expression level between relocations, similar to inversion (See Fig. 1). Adjusted p-values are computed using the Wald-test and Benjamini-Hochberg multiple test correction as implemented by the Deseq2 tool where n.s. indicates a non-significant difference, *= padj < 0.01, **=padj < 0.001 (n=2) (**Supplementary Table S2**).

Similar to *Pitx1^EGFP;Inv1+/-^*, *Pitx1^EGFP;ΔPen^*^;*Rel1+/-*^ E12.5 forelimbs showed 2% EGFP+ cells, suggesting that at this location the inversion and relocations bear a similarly mild transcriptional effect on *Pitx1* and the EGFP sensor (Fig. 2D, E). In contrast, in *Pitx1^EGFP;ΔPen^*^;*Rel2+/-*^, we measured 59% of EGFP+ forelimb cells (Fig. 2D, E), two times more as the one observed when placing *Pen* at the same position in *Pitx1^EGFP;Inv2+/-^* forelimbs (27% see Fig.1E). This difference suggests that the alterations in CTCF relative positioning and binding site directionality within the inverted interval might restrict the capacity of *Pen* to induce *Pitx1* in *Pitx1^EGFP;Inv1+/-^* forelimbs. Indeed, in contrast to *Pitx1^EGFP;ΔPen^*^;*Rel2+/-*^, the *Pitx1^EGFP;Inv2+/-^* allele causes the relocation and inversion of a *Pitx1*-convergent CTCF binding site at *PDE*, to the telomeric inversion breakpoint (**Supplementary Fig. S2**). Finally, in the most proximal relocation, *Pitx1^EGFP;ΔPen^*^;*Rel3+/-*^, 62% of E12.5 forelimb were found expressing EGFP (Fig. 2D, E). Overall, the similar proportion of EGFP+ cells in *Pitx1^EGFP;ΔPen^*^;*Rel2+/-*^ and *Pitx1^EGFP;ΔPen^*^;*Rel3+/-*^, shows that repositioning the enhancer either in the *PDE* region or a few kb upstream of the gene promoter induces a similar effect on *Pitx1* mis-activation (Fig. 2D, E**, Supplementary Fig. S2**).

As inversions showed a similar transcription level between alleles in EGFP+ cells, we wanted to confirm this in the context of the relocations. We therefore performed RNA-seq in EGFP+ cells and found that *Pitx1* and EGFP expression is similar in all the active cells (Fig. 2F**, Supplementary Table S2**). Overall, this data shows that the ability of *Pen* to contact *Pitx1* defines the proportion of cells in which the gene will be ectopically activated, yet, it does not strongly affect *Pitx1* transcription level per allele.

### Increase in *Pitx1* ectopically expressing forelimb cells associate with worsened skeletal defects

As changes in *Pen* positioning lead to a different proportion of cells ectopically activating *Pitx1*, the phenotypic effect of these variations is unknown. To test whether an increase in affected cells is linked to a worsened phenotype, we analysed mutant skeletons of E18.5 embryos and scored forelimb malformations. We decided to compare wildtype to *Pitx1^EGFP;Inv1+/-^*, *Pitx1^EGFP;Inv2+/-^* and *Pitx1^EGFP;ΔPen^*^;*Rel3+/-*^ skeletons as these three precisely showed a progressive increase in EGFP+ cell proportions with 6.4%, 27%, and 62%, respectively. Weakly overexpressing forelimbs from *Pitx1^EGFP;Inv1+/-^*resulted in a mild phenotype, specifically with a slight bowing of the radius and ulna (Fig. 3A, B**, Supplementary table S3**). Notably, the same allele showed a stronger phenotype when bred to homozygosity and assayed in adult mice (Kragesteen, et al, 2018). *Pitx1^EGFP;Inv2+/-^* forelimbs, where 27% of cells are EGFP+ at E12.5, showed more striking bowing of the radius and ulna (Fig. 3C**, Supplementary Table S1, S3**). Furthermore, we noted a significant reduction of the deltoid crest, a characteristic structure of the forelimb, accompanied by a mildly hypoplastic olecranon. Additionally, there was a noticeable broadening of the distal head of the humerus and the proximal head of the radius, a phenotype that aligns with previous descriptions in patients (Fig. 3C**, Supplementary Table S1, S3**). Finally, *Pitx1^EGFP;ΔPen^*^;*Rel3+/-*^ forelimbs, where 62% of cells are EGFP+ at E12.5, exhibited the most severe phenotype. This included the recurring bowing of the long zeugopodal bones, strong reduction of the deltoid crest, broadening of the distal humerus and proximal radius and notably, in all analysed *Pitx1^EGFP;ΔPen^*^;*Rel3+/-*^ skeletons, an aplastic or severely hypoplastic olecranon, a feature not observed in other alleles, but often Liebenberg syndrome patients (Fig. 3D, **Supplementary table S1, S3**). Finally, we observed a relative thinning of the ulna compared to its radius counterpart, in a similar way as the fibula is thinner than the tibia, underlining the arm-to-leg transformation. Overall, our analysis shows that an increase of *Pitx1* ectopically activating cells has a positive correlation with the accumulation of defects in the developing forelimb skeleton.

**Figure 3:**
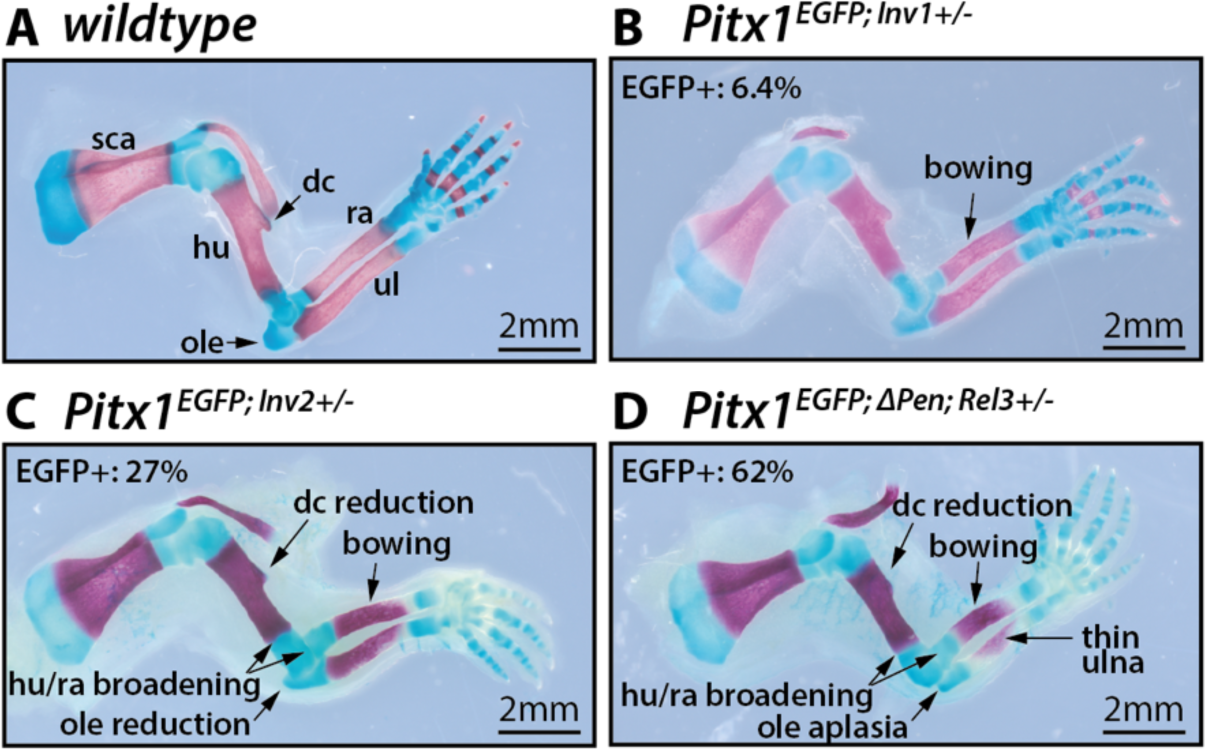
Increasing proportions of Pitx1 ectopically-expressing cells correlates with severity of skeletal defects. **A** Alizarin red and alcian blue staining of wildtype E18.5 forelimbs. Black arrows pinpoint to sca: scapula, hu: humerus, dc: deltoid crest, ole: olecranon, ra: radius, ul: ulna. **B-D**. Alizarin red and alcian blue staining of mutants (**B**) Pitx1^EGFP;Inv1+/-^, (**C**) Pitx1^EGFP;Inv2+/-^ and (**D**) Pitx1^EGFP;ΔPen;Rel3+/-^E18.5 forelimbs. Arrows point to dysplastic skeletal features: bowing of the radius and ulna, reduction of the deltoid crest, reduction of the olecranon, broadening of distal humerus and proximal radius and ulna, relative thinning of ulna.

### *Pitx1* forelimb *endo*-activation mirrors hindlimb *Pitx1* expression

Liebenberg-associated SVs have been described to lead to arms assuming various skeletal and soft tissue features of legs (DeLaurier et al., 2006; Kragesteen et al., 2018; Spielmann et al., 2012). To understand to what extent SV-induced *Pitx1* forelimb transcription resembles its normal hindlimb activity, we performed 10X single-cell RNA-seq (scRNA-seq) on stage-matched E12.5 *Pitx1^Inv1+/-^* forelimbs and compared to wildtype fore- and hindlimbs (Rouco et al., 2021). The first level of clustering revealed six main limb clusters: muscle, neuron, immune cells, epithelium, endothelium and mesenchyme (Fig. 4A) (**Supplementary Table S4**). We noticed that *Pitx1* expression was restricted to the mesenchyme in wildtype hindlimbs but also in *Pitx1^Inv1+/-^* forelimb, although at lower expression levels (**Supplementary Fig. S3A and S3B**). We then subclustered the mesenchyme to obtain more definition to quantify *Pitx1* expression across sub-populations (Fig. 4A). Here, we identified nine mesenchymal populations comparable to the one previously characterised in E12.5 limb mesenchyme (Rouco et al., 2021). Four clusters showed proximal identity: Proximal Proliferative Progenitors (**PPP**), Tendon Progenitors (**TP**), Irregular Connective Tissue (**ICT**) and Proximal Condensations (**PC**). An additional four clusters showed distal identity: Distal Proliferative Progenitors (**DPP**), Distal Progenitors (**DP**), Early Digit Condensations (**EDC**) and Late Digit Condensations (**LDC**). Finally, we identified a Mesopodium (**MS**) cell cluster, neither proximal nor distal. In wildtype hindlimbs and *Pitx1^Inv1+/-^* forelimbs, we observed *Pitx1* expression in all these clusters showing that the forelimb gain of expression occurred with a similar specificity than in hindlimbs. However, the variation in *Pitx1* expression between clusters was more pronounced in wildtype hindlimbs compared to *Pitx1^Inv1+/-^*forelimbs (Fig. 4B, **Supplementary Table S4**). This observation indicates that the mesenchymal specificity of *Pitx1* expression is preserved in mutant forelimbs when compared to wildtype hindlimbs, albeit not to its full extent across mesenchymal subpopulations.

**Figure 4:**
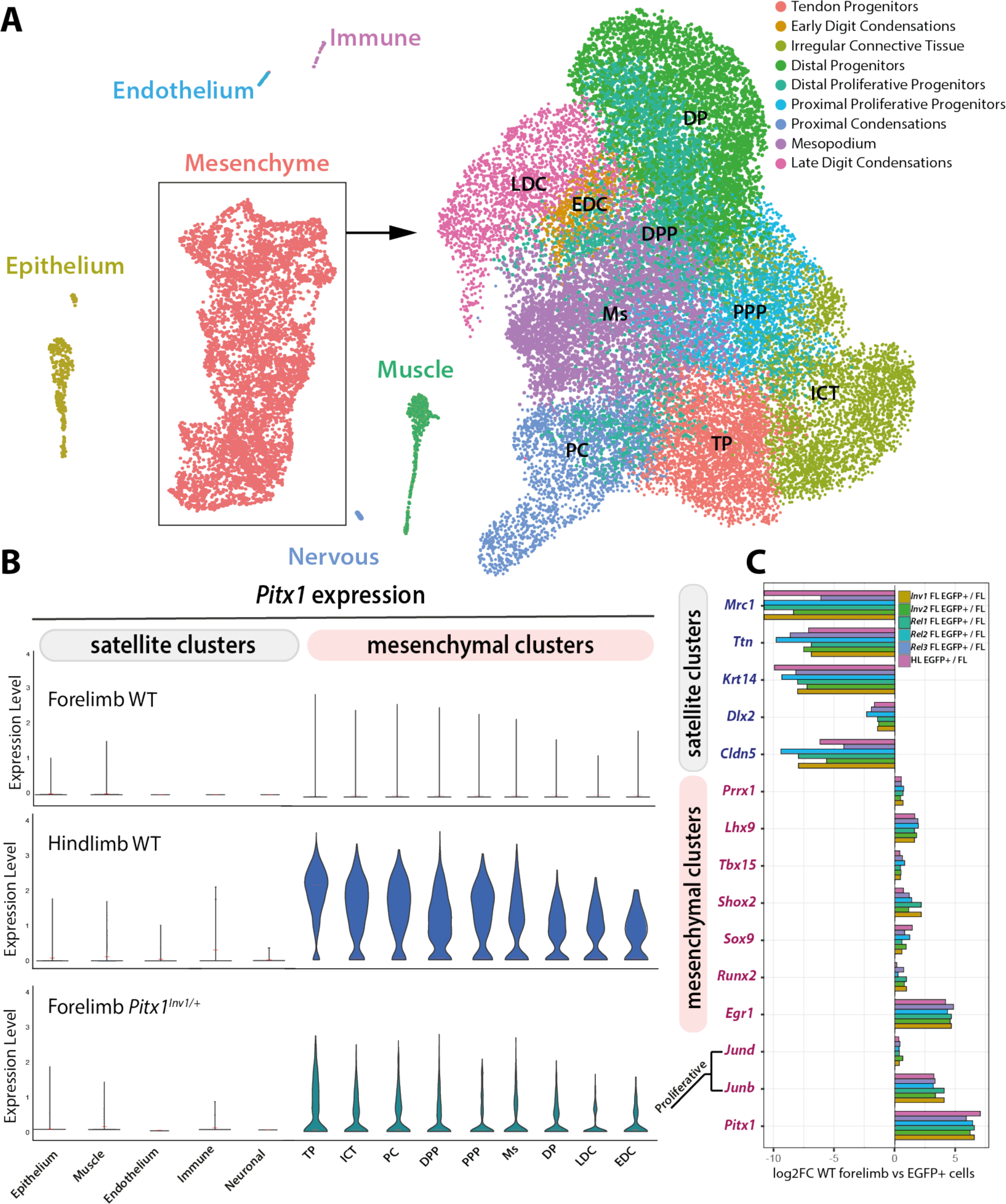
Pitx1 forelimbs ectopic expression reflects the gene’s transcriptional program in hindlimbs. **A.** UMAP of cell clusters present in wildtype fore- and hindlimbs and Pitx1^Inv1+/-^ hindlimbs: right all cells, left mesenchymal cell type sub-clustering. **B.** Pitx1 expression by cell cluster in wildtype fore- and hindlimbs and Pitx1^Inv1+/-^ forelimbs. Not the overall similarly of expression between wildtype hindlimbs and Pitx1^Inv1+/-^ forelimbs. **C.** Selected marker genes enrichment across EGFP+ population of Pitx1^EGFP^ hindlimbs, as well as Pitx1^EGFP;Inv1+/-^, Pitx1^EGFP;Inv2+/-^, Pitx1^EGFP;ΔPen;Rel1+/-^, Pitx1^EGFP;ΔPen;Rel2+/-^, Pitx1^EGFP;ΔPen;Rel3+/-^ forelimb compared to wildtype bulk forelimbs.

To assay whether these expression specificities are a general feature of *Pitx1 endo*-activation, we analysed the enrichment of marker genes in *Pitx1^EGFP;Inv1+/-^*, *Pitx1^EGFP;Inv2+/-^*, *Pitx1^EGFP;ΔPen^*^;*Rel1+/-*^*, Pitx1^EGFP;ΔPen^*^;*Rel2+/-*^, *Pitx1^EGFP;ΔPen^*^;*Rel3+/-*^ forelimbs and control *Pitx1^EGFP^* hindlimb EGFP+ cells compared to wildtype forelimbs. Generally, we observed homogenous marker gene enrichment among mutants, showing high similarity between EGFP+ cells (**Supplementary Table S2**). More specifically, all the EGFP+ populations showed a depletion of genes linked to non-mesenchymal cell identity (*Wnt6, Ttn, Krt14, Dlk2, Cldn5*) and an enrichment for mesenchymal markers (*Prrx1, Lhx9*) confirming that *Pitx1 endo*-activation specifically occurs in mesenchymal cell types (Fig. 4C). We also observed enrichment for proximal (*Shox2* and *Tbx15*), tendons (*Egr1*) and chondrogenic markers *(Sox9, Runx2*) corroborating the previous findings obtained from scRNA-seq. Furthermore, we also found that cell expressing *Pitx1* were enriched for cell division markers as *JunB* and *JunD* in line with the tissue outgrowth properties associated to *Pitx1* (Duboc and Logan, 2011; Rouco et al., 2021) (Fig. 4C). This shows the cell-specificity of *Pitx1 endo*-activation in forelimbs mirrors to a certain extent its physiological expression in wildtype hindlimb.

Finally, to understand whether *Pitx1 endo*-activation can induce a wider hindlimb-like transcriptional program, we compared bulk *Pitx1^EGFP;ΔPen^*^;*Rel2+/-*^ and *Pitx1^EGFP;ΔPen^*^;*Rel3+/-*^ to wildtype forelimbs transcriptome. Here, we detected that the hindlimb-specific gene *Tbx4* was upregulated in mutant forelimbs, indicating that *Pitx1* expression could induce its transcription (**Supplementary Table S5**) (Logan and Tabin, 1999). We also noted an increase in cartilage and chondrogenesis related markers such as *Sox9*, *Foxc1 and Gdf5* suggesting an increased chondrogenic program in mutant forelimbs (**Supplementary Table S5**) (Nemec et al., 2017). Altogether, these findings underline that *Pitx1 endo*-activation establishes, in the forelimb counterpart of hindlimb *Pitx1* expressing cell-types, features of hindlimb transcriptional programs.

### SV-induced *Pitx1 endo*-activation promotes topological changes

Hindlimb cells transcriptionally active for *Pitx1* adopt a fundamentally different 3D locus topology than their inactive counterparts (Rouco et al., 2021). Consequently, it is plausible that SVs-induced *Pitx1 endo*-activation leads to topological changes in transcriptionally active cells. To test this hypothesis, we initially performed C-HiC on *Pitx1^EGFP;Inv1+/-^*, comparing EGFP+ and EGFP-forelimb cells. We found that *Pitx1* contacts *Pen* as well as *PelB*, *PDE*, and *RA4* more frequently in EGFP+ cells than in EGFP-cells (Fig. 5A). Conversely, in EGFP-cells, the repressive contact between *Pitx1* and *Neurog1* was more prevalent than in EGFP+ cells (Fig. 5A). These differences are strikingly similar to those observed between hindlimb EGFP+ and EGFP-cells (Rouco et al, 2021), indicating that this inversion facilitates the formation of an active topology specifically in transcriptionally active cells.

**Figure 5:**
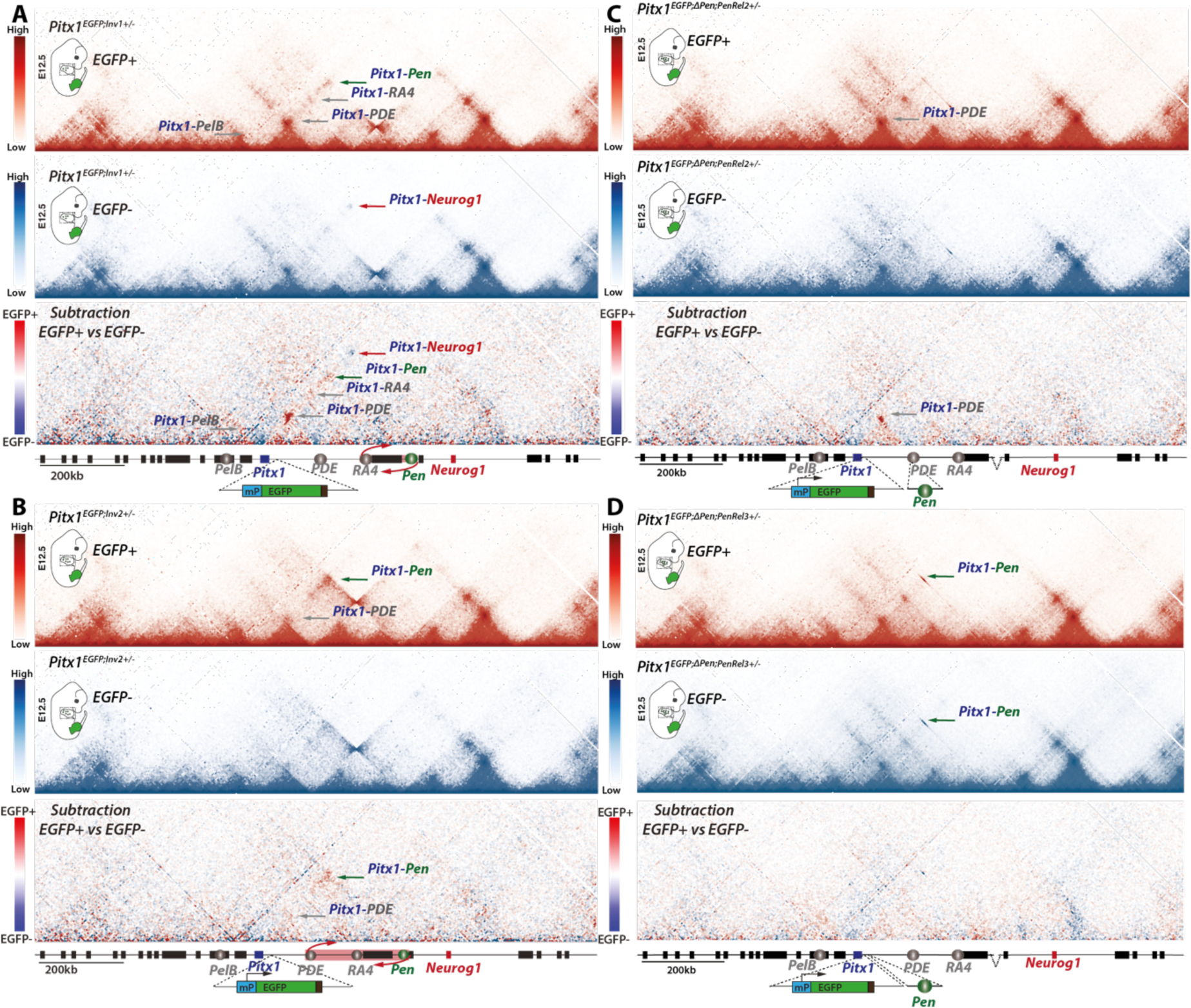
Topological changes at the locus diminish as Pitx1-Pen contact probabilities increase. **A-D** C-HiC of the Pitx1 locus in EGFP+ (red maps) and EGFP-(blue maps) cells from (**A**) Pitx1^EGFP;Inv1+/-^ forelimbs, (**B**) Pitx1^EGFP;Inv2+/-^ forelimbs, (**C**) Pitx1^EGFP;ΔPen;Rel2+/-^ forelimbs and (**D**) Pitx1^EGFP;ΔPen;Rel3+/-^ forelimbs. Darker red or blue bins indicate stronger interaction frequencies as shown on the scale bars. For each panel, the lowest map is a subtraction of the two above where preferential interactions in EGFP+ cells are shown in red, while the ones in EGFP-cells are shown in blue. Contacts between Pitx1 and Pen are shown with a green arrow, Pitx1 contacts with PelB, PDE or RA4 are shown with a grey arrow, the Pitx1-Neurog1 contact is shown with a red arrow. All subtraction scales were homogenized for comparison purposes.

We next investigated whether different active-inactive topologies would also be present in the other alleles described or if this was a specific feature of *Pitx1^EGFP;Inv1+/-^* forelimbs. Thus, we generated C-HiC maps of EGFP+ and EGFP-cells obtained from *Pitx1^EGFP;Inv2+/-^*, *Pitx1^EGFP;ΔPen^*^;*Rel2+/-*^ and *Pitx1^EGFP;ΔPen^*^;*Rel3+/-*^ forelimbs. In *Pitx1^EGFP;Inv2+/-^*, despite a higher proportion of *Pitx1*-expressing cells (See Fig. 1D), we observed fewer changes in interaction between EGFP+ and EGFP-cells. Here, only the interaction between *Pitx1* and *Pen* was strongly increased in EGFP+ cells and, to a lesser extent, that between *Pitx1* and *PDE* (Fig. 5B). Similarly, in *Pitx1^EGFP;ΔPen^*^;*Rel2+/-*^, EGFP+ cells showed a clear gain of contacts between *Pitx1* and *PDE*, where the *Pen* enhancer is relocated, but not with other regions (Fig. 5C). These results consistently highlight strengthened *Pitx1*-*Pen* contact in transcriptionally active cells, suggesting that increased physical proximity is essential for transcription. Lastly, *Pitx1^EGFP;ΔPen^*^;*Rel3+/-*^ EGFP+ and EGFP-forelimb cells exhibited limited topological changes (Fig. 5D). Here, due to the short 7.7kb interval between *Pitx1* and *Pen*, the contact frequency between the two elements was very high in both active and inactive cells. Yet, we noted a relatively stronger contacts in EGFP-cells, a phenomenon already observed for active short-range regulatory contact (Fig. 5D) (Benabdallah et al., 2019). In conclusion, across the different gain-of-function alleles, we observe that fewer locus-wide topological changes are linked to activation when *Pen* is closest to *Pitx1*.

We further explored whether changes in chromatin topology are associated with changes in cis-regulatory element activities by performing H3K27ac Chromatin Immunoprecipitation (ChIP-seq) in *Pitx1^EGFP;Inv1+/-^*, *Pitx1^EGFP;ΔPen^*^;*Rel2+/-*^ and *Pitx1^EGFP;ΔPen^*^;*Rel3+/-*^ EGFP+ cells. As expected, we observed a strong enrichment of H3K27ac at the *Pitx1* promoter in all cases. Moreover, in both *Pitx1^EGFP;Inv1+/-^* and *Pitx1^EGFP;ΔPen^*^;*Rel2+/-*^, there was an increase in H3K27ac coverage at *PDE*, a region interacting with *Pitx1* in both alleles. Finally, in *Pitx1^EGFP;ΔPen^*^;*Rel3+/-*^ EGFP+ cells, only the region adjacent to the *Pen* relocation showed a clear acetylation signal (**Supplementary Figure S4**). We also noted that in the two relocation alleles, the loss of *Pen* at its endogenous genomic location resulted in decreased H3K27ac spreading around it, while an increase around the *Pen*-relocated region was observed, showcasing the spreading potential of the histone mark. In summary, the increased chromatin contacts observed in C-HiC data always involved regions marked by H3K27ac.

### Targeted activation of *Pitx1* does not induce topological change

In the context of SV-induced *Pitx1 endo*-activation, the relocation of the *Pen* enhancer associates with changes in transcriptional activity and genome topology (See Fig. 5 and **Supplementary Fig. S4**). Because both events occur in the same cells, it is unclear whether it is the transcription of the locus that induces the 3D topological changes or whether these occur independently. To assay whether ectopic activation of *Pitx1* is sufficient to induce changes in the locus topology, we first developed an *in vivo* dCas9-P300 activator targeted to the *Pitx1* promoter. To achieve specific expression of the activator in cell clusters permissive to *Pitx1* expression (See Fig. 5), we integrated the dCas9-P300 transgene preceded by a minimal promoter, as a sensor, upstream of the *Shox2* gene promoter to produce *Shox2^dCas9P300/+^* mESCs (Fig. 6A). We selected *Shox2* because of its similar expression specificity with *Pitx1* in developing hindlimbs (correlation coefficient=0.577, p-value = 0.001, where the p-value is the probability for the correlation coefficient to be negative). To direct the dCas9-P300 activator to *Pitx1*, we integrated two sgRNAs that target the *Pitx1* transcriptional start site (TSS) at the *ColA1* locus to produce *Shox2^dCas9P300/+^;ColA1^TSSsgR^*ESCs (Fig. 6A) (Beard et al., 2006).

**Figure 6:**
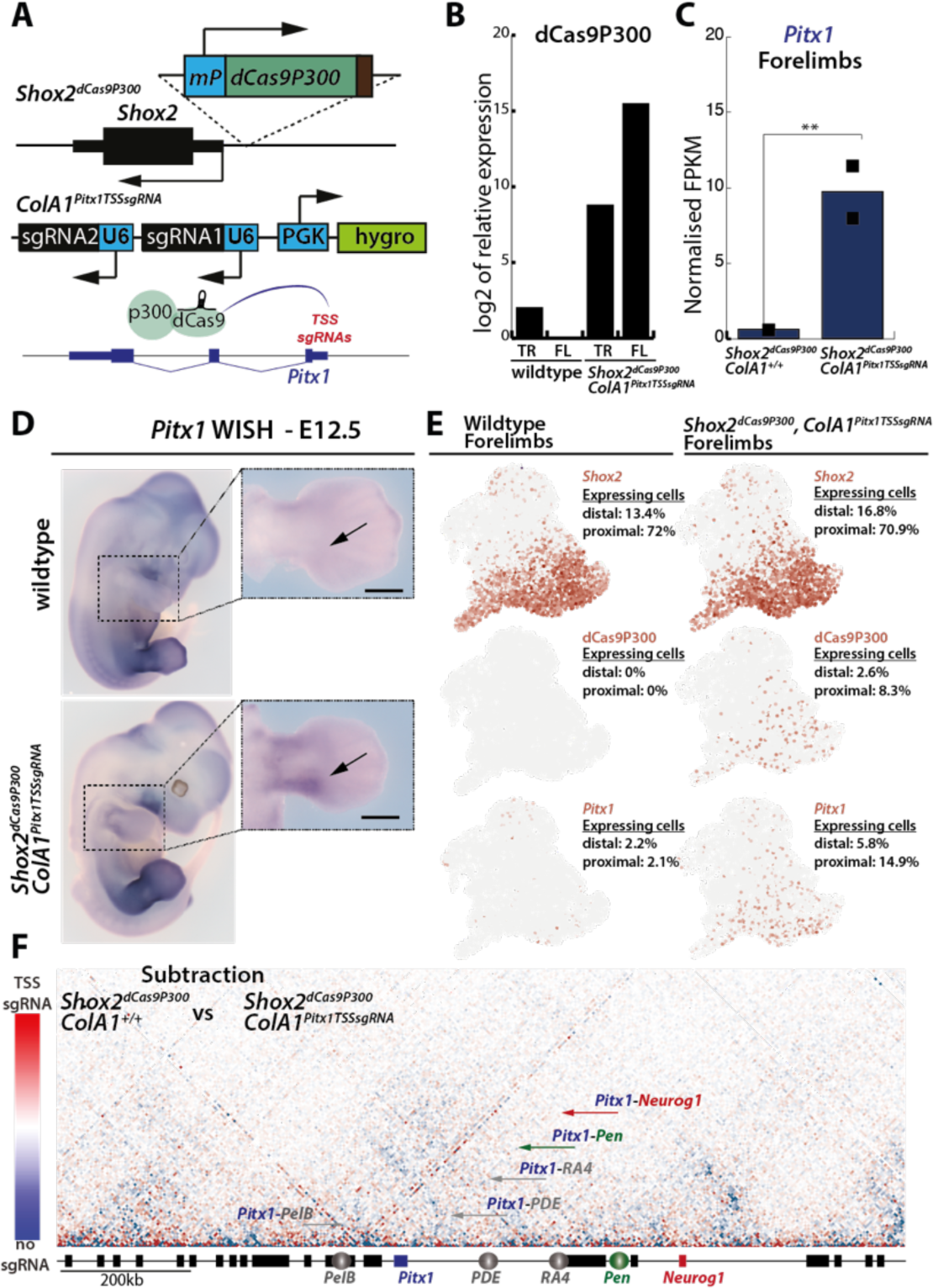
dCas9P300 induces Pitx1 expression in forelimbs without topological changes. **A**. A dCas9-P300 cassette was inserted as a sensor upstream of the Shox2 promoter, two sgRNAs to target dCas9 activity were integrated at the ColA1 safe harbour locus through an FRT-mediated recombination. **B**. RT-qPCR of dCas9P300 in wildtype and Shox2^dCas9P300/+^;ColA1^TSSsgR^ E12.5 forelimbs (FL) and trunk (TR) tissues. The values represent a log2 fold change compared to wildtype forelimb that was set to 1. **C**. Normalised Pitx1 FPKMs in Shox2^dCas9P300/+^;ColA1^+/+^ and Shox2^dCas9P300/+^;ColA1^TSSsgR^. Adjusted p-values are computed using the Wald-test and Benjamini-Hochberg multiple test correction as implemented by the Deseq2 tool, where **=padj < 0.001 (n=2) (**Supplementary Table S6**). **D**. WISH of Pitx1 in wildtype and Shox2^dCas9P300/+^;ColA1^TSSsgR^ forelimbs. Note the proximal gain of Pitx1 expression (black arrow). **E**. Individual UMAPS of scRNA-seq data from wildtype and Shox2^dCas9P300/+^;ColA1^TSSsgR^ forelimbs showing the distribution of Shox2, Pitx1 and dCas9P300 expressing cells as well as the respective percentage of expressing cells in proximal forelimb (proximal) and distal forelimb (distal). **F**. Subtraction of Shox2^dCas9P300/+^;ColA1^TSSsgR^ and Shox2^dCas9P300/+^;ColA1^+/+^ Pitx1 E12.5 proximal forelimbs C-HiC maps. Contacts more frequent in TSS sgRNA are in colored in red, and those more frequent in no sgRNA are colored in blue (See scale bar on the left). Green arrow points at Pitx1 and Pen contacts, Pitx1 contacts with PelB, PDE or RA4 are shown with a grey arrow, Pitx1-Neurog1 contact is shown with a red arrow. Note the absence of visible change. Corresponding C-HiC maps are shown in **Supplementary Figure S6**.

We then derived E12.5 *Shox2^dCas9P300/+^;Cola1^TSSsgR^*embryos using tetraploid aggregation (Artus and Hadjantonakis, 2011). We could detect dCas9-P300 transcripts in forelimbs but not in the embryonic trunk, confirming the expression specificity of the sensor (Fig. 6B). Using RNA-seq, we measured *Pitx1* expression in *Shox2^dCas9P300/+^;ColA1^+/+^* and *Shox2^dCas9P300/+^;ColA1^TSSsgR^* forelimbs and could detect a 15-fold upregulation of the gene in the latter (Fig. 6C**, Supplementary Table S6)**. As observed by whole mount in-situ hybridization (WISH), the expression pattern of *Pitx1* was localized to the proximal forelimb and reminiscent of *Shox2* expression in E12.5 forelimbs (Fig. 6D). Single-cell RNA-seq revealed that *Pitx1* was expressed in 9% of *Shox2^dCas9P300/+^;ColA1^TSSsgR^* forelimb mesenchyme compared to 2% of wildtype counterparts (Rouco et al., 2021). Moreover, we could generally observe that, *Pitx1* and *Shox2* expression domains colocalized in proximal clusters (*Pitx1*-*Shox2* correlation in the entire *Shox2^dCas9P300/+^;ColA1^TSSsgR^* forelimb=0.441 p-value=0.0005, where the p-value is the probability for the correlation coefficient to be negative Fig. 6E**, Supplementary Figure S5**).

We next tested whether the gain of *Pitx1* transcription would elicit a change in 3D conformation of the locus. To enriched for *Pitx1* transcriptionally active cells, we micro-dissected E12.5 proximal forelimbs of *Shox2^dCas9P300/+^;ColA1^TSSsgR^* (14.9% of *Pitx1*-expressing cells) and *Shox2^dCas9P300/+^;ColA1^+/+^* (2.1% of *Pitx1*-expressing cells) and performed C-HiC (**Supplementary Figure S6**). When compared to *Shox2^dCas9P300/+^;ColA1^+/+^* proximal E12.5 forelimbs we did not observed any changes in locus interactions between the two alleles (Fig. 6F), suggesting that major topological contacts with *PelB, PDE, RA4* and *Pen* are not induced by direct activation of *Pitx1*.

### Loss of PRC2 repression induces *Pitx1* forelimb transcription without topological changes

Because the targeted activation of *Pitx1* affected a limited proportion of forelimb cells, subtle topological changes could be missed. Therefore, we used a different approach to activate *Pitx1* transcription and asked whether removal of PRC2-mediated polycomb repression would be a more effective method. PRC2 is a multiprotein complex made of several subunits including the H3K27me3 reader EED which enables the spreading of the mark over chromatin domains (Piunti and Shilatifard, 2016). Here we exploited a conditional *Eed* floxed allele combined to a full *Eed* knock out, and a limb-specific mesenchymal CRE driver (*Prx1-CRE;Eed^flox/-^*) to assess the effect of its loss on both *Pitx1* transcription and locus structure (Gentile et al., 2019; Logan et al., 2002; Yu et al., 2009).

Through WISH, we could observe a strong gain of *Pitx1* expression in proximal *Prx1-CRE;Eed^flox/-^* E12.5 forelimbs (Fig. 7A). We then re-analysed RNAseq data and observed a 27-fold upregulation of *Pitx1* in mutant forelimbs compare to wildtype littermates (Fig. 7B)(Gentile et al., 2019). As expected, in proximal E12.5 forelimb, a decrease in H3K27me3 could be detected throughout the locus (Fig. 7C) (Guerard-Millet et al., 2021). It is also interesting to note that despite loss of H3K27me3 at *Neurog1*, the gene, unlike *Pitx1*, was not ectopically transcribed in forelimb cells, underlying cell-specificity as a requirement for mis-activation of genes (Fig. 7C). The decrease of H3K27me3 at *Pitx1* also coincided with the accumulation of the active H3K27ac mark at the gene promoter and *PDE* (Fig. 7C, arrows) (Gentile et al, 2019)(Guerard-Millet et al., 2021). This shows that, at *Pitx1,* the removal of PRC2 repression results in the activation of the locus.

**Figure 7:**
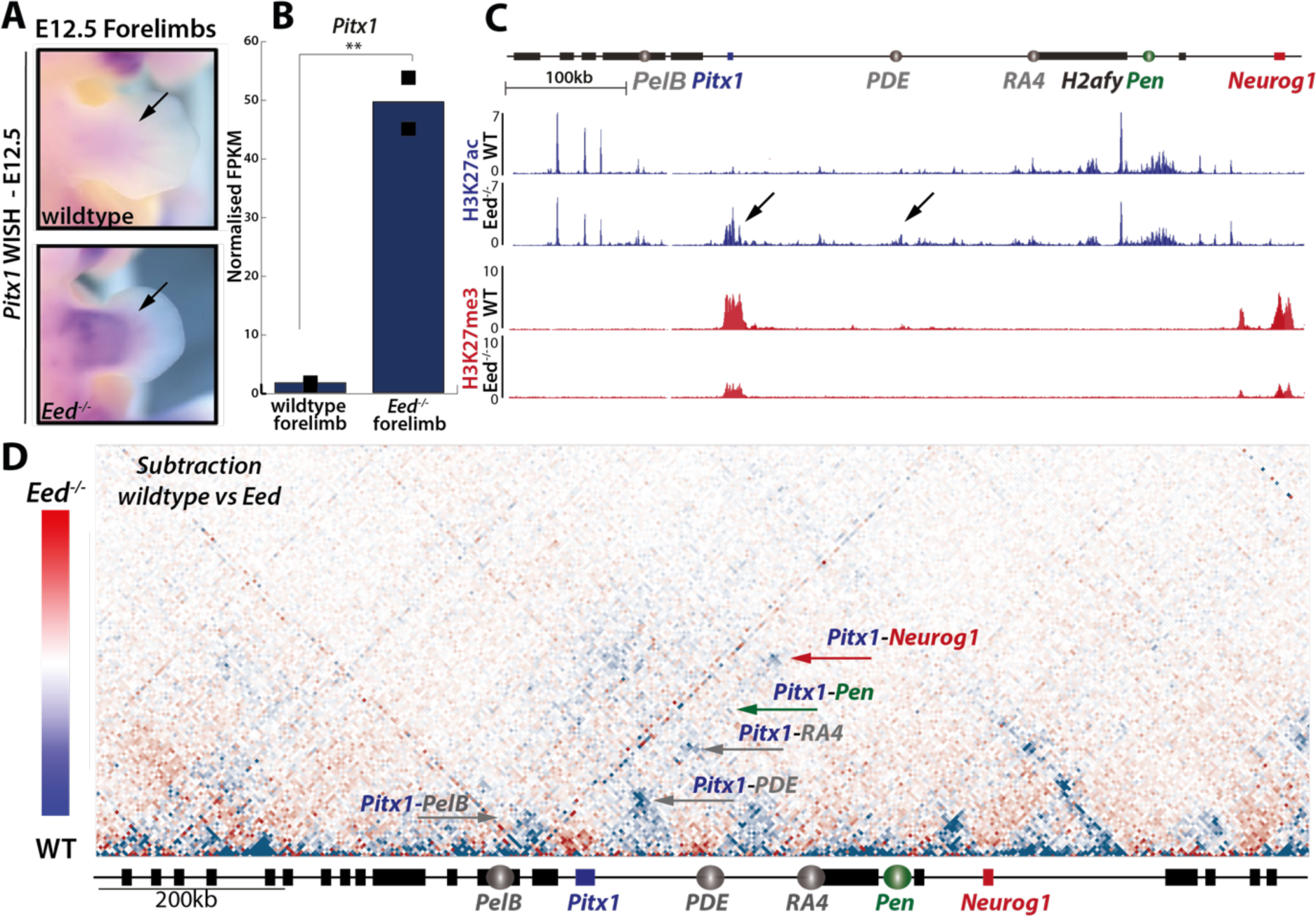
Loss of PRC2 repression leads to Pitx1 expression in forelimbs. **A**. Pitx1 WISH of E11.5 wildtype and Prx1-Cre;Eed^flox/-^ (Eed^-/-^) forelimbs. Note the strong proximal gain of Pitx1 expression (black arrow). **B**. Normalised FPKMs of Pitx1 in E12.5 wildtype and Prx1-CRE, Eed^flox/-^ (Eed) forelimbs. Adjusted p-values are computed using the Wald-test and Benjamini-Hochberg multiple test correction as implemented by the Deseq2 tool, tool where **=padj < 0.001 (n=2). **C**. ChIP-seq of H3K27ac (first two tracks) and H3K27me3 (second set of tracks) show an accumulation of H3K27ac at the Pitx1 locus (black arrows) in proximal Prx1-Cre;Eed^flox/-^ (Eed^-/-^) compared to wildtype (WT) forelimbs and an overall reduction of H3K27me3 signal. **D**. Subtraction of wildtype and Prx1-Cre;Eed^flox/-^ (Eed) E12.5 proximal forelimbs C-HiC maps. Contacts more frequent in Eed are colored in red, and those more frequent in widtype are colored in blue (See scale bar on the left). Green arrow points at Pitx1 and Pen contacts, Pitx1 contacts with PelB, PDE or RA4 are shown with a grey arrow, Pitx1-Neurog1 contact is shown with a red arrow. Corresponding C-HiC maps are shown in **Supplementary Figure S7**.

We then explored whether the loss of PRC2 repression results in a change of topological organisation of the locus (**Supplementary Figure S7**). First, we observed a reduction of the *Pitx1*-*Neurog1* PRC2-associated contact in *Prx1-Cre;Eed^flox/-^* proximal forelimbs compared to wildtype (Fig. 7D). However, similarly to the dCas9-P300 C-HiC data, we did not observed a gain of interactions between *Pitx1* and its enhancers in *Prx1-Cre;Eed^flox/-^* forelimbs (Fig. 7D). In fact, we observed a relative loss of the contacts with *PelB, PDE, RA4* and *Pen*, which suggests that PRC2 loss leads to a disorganisation of the locus topology. We concluded that loss of PRC2 leads to *Pitx1* activation independently from strengthening of enhancer-promoter topological contacts and therefore that transcription is not sufficient to induce changes in locus topology at the *Pitx1* locus.

## Discussion

In this work, we show that changes in the relative positioning between *Pitx1* and its *Pen* enhancer associate with a variable proportion of *Pitx1* overexpressing cells in developing forelimbs. Within this active cell population, the levels of *Pitx1* expression do not increase with enhancer-promoter proximity but rather reach a conserved threshold of activation. This suggests that once activation is achieved at the *Pitx1* locus it is done so at its full transcriptional potential where *Pitx1* promoter activity is saturated.

Changes in *Pitx1*-*Pen* distance and its associated variation in the proportion of cells ectopically expressing *Pitx1*, but not in *Pitx1* transcription per allele, provides a mechanistic framework to account for the variation in Liebenberg syndrome severity among cases described so far. Here, we have shown that the more a SV reduces the *Pen*-*Pitx1* distance, and consequently the number of intermediate CTCF sites, the higher the proportion of forelimb *Pitx1* overexpressing cells will be and the stronger the skeletal defects. Similarly, patients with SVs inducing a short genomic distance and few intermediate CTCF binding between *Pitx1* and *Pen* displayed more severe malformations (**Supplementary Fig. S1**). In general, variability in rare disease severity was already described in several cases. For instance, several overlapping deletions at the *Epha4* locus, that induce rewiring of enhancers toward the *Pax3* gene, result in brachydactyly and variable hand defects (Lupianez et al., 2015). Here, the proportion of cells affected by the *Pax3* overexpression in the distinct SVs could explain the variability in phenotypical outcome. In another reported case, different duplications at the *Ihh* locus, leading to variable increases of gene expression in developing limbs, were also shown to result in variable syndactyly phenotypes. Moreover, LacZ analysis of *Ihh* in the mutants indicated broadened expression domains of the gene, suggesting that increase in expression could be due to more cells ectopically activating *Ihh* (Will et al., 2017). Therefore, although our data provides a mechanism for variation in the Liebenberg syndrome, it could be applied to many other syndromes linked to ectopic gene transcription.

In a previous study, we have shown that the homozygous loss of *Pen* did not result in a full *Pitx1* loss-of-function in hindlimbs, but in a 30% reduction of *Pitx1* transcription (Rouco et al, 2021). This hindlimb loss was mostly the result of a fraction of cells from all mesenchymal clusters, without further specificity, not displaying any *Pitx1* transcription. It was therefore hypothesised that *Pen* acts as a “support” enhancer enabling the robust *Pitx1* transcriptional initiation in the mesenchyme. In this perspective, other regions would act to provide more cell-type specificity, such as RA4 that was recently described as a chondrogenic enhancer (Darbellay et al., 2023). This is similar to what happens during *endo*-activation, where *Pen* activates *Pitx1* in all forelimb mesenchymal clusters without further specificity. Together, *Pen*-dependent loss and gain of *Pitx1* expression pinpoint to the same role for *Pen*: to act as a *pan*-mesenchymal enhancer with the ability to trigger robust transcriptional onset at the *Pitx1* locus. As in hindlimbs other enhancers are required to further define *Pitx1* cell-type specific expression, it remains to be shown, whether other local enhancers, such as *RA4* which is also active in forelimbs, contribute to the final *Pitx1 endo*-activated expression in forelimbs.

By comparing the locus 3D topology in active and inactive cells, we observed that alleles driving *Pitx1* expression in a limited proportion of cells displayed the most extensive topological changes. Specifically, in the smallest inversion, *Pitx1^EGFP;Inv1+/-^*, multiple enhancer-promoter contact are observed in active cells similar to the previously described stack configuration that occur, in fact, only in a fraction of *Pitx1*-expressing hindlimbs (Hung et al., 2024). In the other extreme, when the *Pen* enhancer was introduced directly upstream of *Pitx1*, in *Pitx1^EGFP;ΔPen;Rel3+/-^,* topologies were very similar between inactive and active cells. Together these data suggest that genetic configurations that reduce the searching space of the *Pitx1* promoter to find *Pen*, i.e. where the *Pen*-*Pitx1* contact is a very probable choice, are more likely to result in expression in a larger proportion of cells. From another perspective, this shows that when transcription is obtained without the need of long-range contacts in the first place, permissive active topologies are not detected. In contrast, in less efficient configurations, i.e. where *Pitx1-Pen* contacts are less likely, active cells display a larger variety of configurations, where *Pitx1* establishes contact patterns with other regulatory regions (*PelB*, *PDE*, *RA4*).

These changes in topology can be the result of two processes: 1) that the transcriptional activation of *Pitx1* increases its ability to form enhancer-promoter contact or 2) that increased enhancer-promoter contacts are required to activate *Pitx1*. Yet, when the *Pitx1* promoter was activated *via* an exogenous dCas9-P300 activator or *via* the alteration of PRC2 activities, we could not observe a gain of enhancer-promoter contact in forelimbs. Despite the caveat of a limited efficiency and thus possible dilution of the signal, the absence of topological effect in the dCas9-P300 targeted *Pitx1* activation is similar to what was observed in the exogenous activation of *Zfp42* by dCas9-VP64 and of *Shh* by TALE-Vp16 in ESCs (Benabdallah et al., 2019; Bonev et al., 2017). In the case of PRC2, as expected from previous data performed on *Eed* knockout mESCs, the repressive contact between *Pitx1* and *Neurog1* was reduced (Denholtz et al., 2013), yet, the activation of *Pitx1* was also not associated to increased enhancer-promoter contacts. Together these data clearly show that transcription, by itself, cannot induce changes in enhancer-promoter contacts at the *Pitx1* locus. It further suggests that changes in enhancer-promoter interactions in a wildtype condition are required, in the first place, to alter the *Pitx1* promoter state *via* its de-repression.

## Material and Methods

### Animal procedures

All animal procedures were in accordance with institutional, state, and government regulations (Canton de Genève authorizations GE/89/19 and GE192A). Animal procedures at the Institut de Recherches Cliniques de Montréal (IRCM) was reviewed and approved by the IRCM animal care committee (protocols 2020-01 and 2021-04).

### Genetically engineered alleles

Engineered alleles using CRISPR/Cas9 technology were created in accordance with the methodology outlined in (Andrey and Spielmann, 2017). sgRNAs were designed using the Benchling software, selecting them based on predicted on-target and off-target scores. Detailed information on all sgRNAs and their corresponding genomic locations for CRISPR–Cas9 can be found in **Supplementary Table S7**. The sgRNAs were sub-cloned into the pX459 plasmid from Addgene, with 8 μg of each vector utilized for the transfection of mESCs. Standard procedures for mESCs culture and genetic editing, were followed. The *Pitx1^GFP^* mESCs clone used was previously described in reference 20. Requests for transgenic G4 ESCs clones can be accommodated.

### Skeletal preparation

Skeletal preparation followed protocols previously described (Paliou et al., 2019). Briefly, sacrificed foetuses were heatshocked at 70°C for 30’’ and skin and viscera were removed. The embryos were fixed in 100% EtOH at room temperature overnight and then stained in Alcian Blue (150 mg/l Alcian Blue 8GX (Sigma-Aldrich) ON at room temperature. Alcian Blue was then washed away with 100% EtOH and replaced with Alzarin Red (50 mg/l Sigma Aldrich) in 0.2% KOH over two days. Finally, the remaining tissue was digested in 1% KOH with visual inspection and skeletons were stored in 0.2%KOH-30% glycerol for imaging and then long-term in 60% glycerol.

### Whole mount *in situ* hybridization

*Pitx1* WISH was performed on E12.5 embryos with a digoxigenin-labelled *Pitx1* antisense probe designed from a cloned antisense probe (PCR DIG Probe Synthesis Kit, Roche 11636090910). Experimental procedure followed the protocol outlined in (Kragesteen et al., 2018).

### Imaging

Embryos were imaged in PBS and skeletons in 0.2%KOH-30% glycerol on an Axio Zoom V16 (ZEISS) microscope. GFP laser exposure was set to 3000 ms.

### Preparation of Single-Cell Limb Suspension

E12.5 limb tissues were microdissected in cold PBS and pooled for processing. To maintain efficiency in downstream experiments, no more than 6 limbs were pooled together at a time. The tissues were dissolved in 400μL Trypsin-EDTA and 40μL 2.5% BSA (Sigma Aldrich, A7906-100G) over 12 minutes at 37°C in a Thermomixer set at 1500 rpm, with a brief resuspension at the 6-minute mark. Trypsin was quenched by adding 400μL 2.5% BSA, and the homogenised tissue was passed through a 40μm cell strainer. An additional volume of 2.5% BSA was passed through to collect any remaining cells. The collected cells were then centrifuged 5’ at 4°C and 400 x g, followed by resuspension in 1% BSA. If H3K27ac ChIP was planned as a downstream experiment, 5mM NaButyrate was added to the 1% BSA.

### Preparation for Single-Cell RNA-seq and Library Construction

Following the preparation of a single-cell limb suspension, cells were counted using an automated counter and resuspended to achieve a concentration of 1400 cells/μL. 50μL of this suspension were provided to the iGE3 Genomic Platform for 10X Library Preparation. The platform performed library preparation for *Pitx1^Inv1+/-^* using the Chromium Single Cell 3’ GEM, Library & Gel Bead Kit v3.0 following the manufacturer’s protocol. Libraries were pair-end sequenced on an Illumina HiSeq 4000 with approximately 8029 cells loaded on a Chromium Chip. For *Shox2^dCas9P300/+^;Cola1^TSSsgR^*library preparation was done using the Chromium Single Cell 3’ GEM, Library & Gel Bead Kit v3.1 following the manufacturer’s protocol. Libraries were pair-end sequenced on an Illumina NovaSeq 6000 with approximately 10,141 cells loaded on a Chromium Chip.

### Cell Sorting

Fluorescence-activated cell sorting (FACS) was employed to identify and sort distinct cell populations in this study, utilizing the Biorad S3 with GFP laser (excitation wavelength 488nm). To eliminate debris from the analysis, FCC/FCS settings were established between 30/40 and 230/220. The viability stain Draq7 was employed to distinguish live cells, and standard protocols were applied to select for singlets. For each sample, a negative control tissue, the embryo’s tails, was included to ensure the purity of the GFP-positive population. Moreover, the gating of GFP-positive populations was consistently applied across multiple experiments to ensure the selection of uniform populations and mitigate variability in GFP intensity over time. FlowJoTM Software was utilized for exporting the analysis in histogram format.

### Cell Processing for ChIP-seq and C-HiC

After sorting, cells were suspended in 1% BSA and then centrifuged 5’ at 400 x g at 4°C in a tabletop centrifuge. The supernatant was discarded, and cells were resuspended in 10% FCS/PBS before fixation at room temperature. For ChIP, 1% formaldehyde was used, and for C-HiC, 2% formaldehyde was applied, both for a duration of 10’ with rolling. Fixation was quenched by adding 1.45M cold glycine, followed by centrifugation at 1000 x g, 8’, 4°C. Cells were then resuspended in cold lysis buffer (10 mM Tris, pH 7.5, 10 mM NaCl, 5 mM MgCl2, 1 mM EGTA, Protease Inhibitor (Roche, 04693159001)). After 10’ of incubation on ice, fixed nuclei were isolated through a 3-minute centrifugation at 1000 x g at 4°C, followed by washing in cold 1 x PBS buffer (1000 x g, at 4°C for 1 minute). The PBS was removed, and nuclei were stored at −80°C.

### Cell Processing for RNA-seq and Library Preparation

For bulk limb analysis, two independent limbs were microdissected and snap-frozen at −80°C for subsequent total RNA extraction using the RNEasy Mini Kit (QIAGEN, 74134) following protocol. RNA quantification was performed with Qubit 2.0 (LifeTechnologies) and the RNA Broad Range Assay (Q10210).

For GFP population studies, after sorting, at least two replicates of 2.5 x 10^5^ cells were pelleted 5’ at 400 x g, 4°C. After removal of 1% BSA, cells were snap-frozen at −80°C for total RNA extraction. RNA extraction was carried out with the RNEasy Micro Kit (QIAGEN, 74004) following the manufacturer’s instructions. Quantification was performed with Qubit and RNA High Sensitivity Assay (Q32852).

Library preparation and sequencing were conducted at the iGE3 Genomic Platform. RNA integrity was assessed with a Bioanalyzer (Agilent Technologies). The SmartSeq v4 kit (Clontech) was used for reverse transcription and cDNA amplification, following the manufacturer’s instructions, with 5ng RNA as input. Library preparation followed with a 200pg cDNA input, using the Nextera XT kit (Illumina). Libraries were assessed by Tapestation and Bioanalyzer with a DNA High Sensitivity Chip, 2nM were pooled and sequenced on an Illumina NovaSeq 6000 sequencer using SBS TruSeq chemistry with an average of 35 million reads (single-end 50bp) per library.

### Chromatin Immunoprecipitation and Library Preparation

For H3K27ac ChIP, an average of 5 x 10^5^ nuclei, and for H3K27me3 ChIP 1 x 10^6^ nuclei were used for each experiment. These were sonicated to an average size of 200-500bp fragments on a Bioruptor Pico Sonicator (Diagenode) for 8 minutes 30 seconds ON/OFF at 4°C. Immunoprecipitation was performed as described previously (Lee, et al., 2006; Jerkovic, et al., 2017), using 3.6μg of chromatin. The antibody used was ⍺-H3K27Ac (Diagenode C15410174) at a 1/500 dilution, 5mM of Na-Bu was added to all buffers.

### Immunoprecipitation

Before sonication, magnetic beads were pre-cleared with 30μL of Protein G beads (for H3K27ac – Invitrogen 10003D) or Protein X beads (for H3K27me3 – Invitrogen 10001D) and 0.25% BSA in PBS. After the addition of the antibody, the beads were left to rotate at 4°C for at least 4 hours. Unbound antibodies were removed, and following sonication, the chromatin was added to fresh sonication buffer and incubated rotating overnight at 4°C. Unbound chromatin was then removed by seven washes in RIPA buffer and one in TE buffer. Chromatin was eluted and de-crosslinked overnight with the addition of 5μL Proteinase K (10mg/mL). RNase A (4μL, 10mg/mL) treatment followed, and then phenol:chloroform:IAA extraction and precipitation. Chromatin was eluted in 50μL H_2_O.

### Library Preparation and Sequencing

Library preparation was performed by the iGE3 Genomic Platform. The Illumina ChIP TruSeq protocol was followed with a <10ng DNA input, and libraries were sequenced as 50bp single-end reads with the Illumina NovaSeq 6000 sequencer. Libraries were validated on Tapestation and Qubit fluorimeter, pooled as 2nM, and sequenced with TruSeq SBS chemistry.

### Capture-HiC and Library Preparation

C-HiC experiments were conducted as singlets using an average of 1×10^6^ fixed nuclei for sorted cells and 3×10^6^ mESC cells. The experiments adhered to the protocol outlined in Kragesteen et al., 2018, and Paliou et al., 2019. In this process, chromatin underwent digestion with the DpnII enzyme (1000U total; NEB, R0543M) at 37°C overnight, supplemented with 20% SDS and 20% Triton X-100. Subsequent ligation was carried out with 100U of ligase in a 1.15% Ligation buffer at 16°C for 4 hours, followed by 30 minutes at room temperature. The decrosslinking step occurred overnight at 65°C with the addition of 30 μL Proteinase K (30mg/mL). RNAse A treatment (30μL, 10mg/mL) was followed by phenol:chloroform:IAA extraction and an overnight precipitation. After precipitation, the DNA pellet was reconstituted in 150μL Tris pH7.5. Total DNA quantification was performed using the Qubit High Sensitivity DNA Assay (Q32851).

### Preparation of 3C Library and Sequencing

Libraries were prepared by the iGE3 Genomic Platform. In brief, chromatin was sheared, and adapters were ligated following the manufacturer’s protocol for Illumina sequencing (Agilent). Libraries underwent pre-amplification and hybridization on custom Sure Select beads spanning the chr13: 54,000,001–57,300,000 region, indexed for sequencing as 50bp paired-end reads (Agilent). Once again, 2nM of libraries were clustered for sequencing on an Illumina Novaseq 6000 with SBS TruSeq chemistry.

### Data Analysis

#### RNA-seq

Reads from RNA-seq were mapped using the STAR 2.7.2b mapper with default settings to the GRCm39/mm39 genome. Output BigWig files were displayed on the UCSC genome browser. Counts were compiled from STAR counts using R 3.6.2, and FPKM were computed through Cufflinks 2.2.1. Normalized FPKM values were calculated by first determining coefficients extrapolated from a set of 1,000 housekeeping genes known for their stable expression as defined from the comparison of a series of RNA-seq (Brawand et al., 2011). The coefficients obtained were then applied to adjust the respective FPKM values. Differential expression analysis utilized the DEseq2 R package, with the Wald test for comparisons across samples and multiple test correction using the FDR/Benjamini-Hochberg test. Each analysis included two biological replicates per condition. Fold-enrichment of *Pitx1* and was calculated using DEseq2’s normalization by size factor.

##### Custom Genomes

For RNA-seq analysis, custom mm39 genomes were generated using STAR 2.7.2b, incorporating an additional chromosome to accommodate the custom sequences of EGFP or dCas9-P300 and polyA tails. The gft file was modified to specify these sequences as coding genes and exons. Cell Ranger 6.1.2 was utilized for single-cell RNA-seq analysis, creating a custom mm39_dCas9P300 genome by adding an extra dCas9P300-containing chromosome and customizing the reference gtf file.

#### ChIP-seq

ChIP-seq reads were mapped to the reference GRCm39/mm39 genome using Bowtie 2.3.4.2 or Bowtie2 2.3.5.1, respectively. Reads were filtered for quality, and BedGraphToBigWig was used to convert files into BigWig format for visualization in the UCSC browser.

#### Capture-HiC

Capture-HiC data analysis followed previous descriptions. Reads were mapped against the reference NCBI37/mm9 genome using Bowtie2 2.3.4.2. Filtering, de-duplication, and processing of valid pairs were done with HiCUP 0.6.1 and Juicer Tools 1.9.9. Binned contact maps were produced with MAPQ ≥30 and exported at 5kb resolution.

#### Single Cell RNA-Seq

Analysis of single-cell RNA-seq involved processing sequenced reads using the 10X Genomics Cell Ranger 6.1.2 software. Data filtering, quality control, normalization, scaling, dimensional reduction, and doublet identification were performed using Seurat 4.3.0 and DoubletFinder 2.0.3.

##### Merging and Normalization

Following individual dataset filtering and normalization, the two wildtype forelimb replicates were merged into a single Seurat object. To account for potential variance due to cell-cycle variations, cell cycle regression was implemented using the CellCycleScoring method with a predetermined list of marker genes (Tirosh et al., 2016). The dataset underwent additional normalization through SCTransform with standard parameters, incorporating the scored cell-cycle and the dCas9P300 feature as regressed variables (Hafemeister et al., 2019).

##### Clustering of Whole Limbs and Mesenchyme

The cells were clustered after cell cycle and dCas9P300 regression using the SCTransform Seurat package. For clustering, PCA (50 npcs) and UMAP (50 dims) were utilized, and the closest neighbors of each cell were calculated. The Seurat FindClusters function was employed with a resolution of 0.1, defining 9 clusters. Cluster identification was performed with the FindMarkers function, enabling the selection of differently expressed gene markers among clusters (ident.1, only.pos=TRUE).

Given the exclusive expression of *Pitx1* and *Shox2* in the mesenchymal cells of the limb, downstream analysis focused on these populations. The 3 mesenchymal cell populations were merged and reclustered. PCA of 20 npcs and UMAP of 20 dims were applied, and closest neighbours were calculated for each cell. Using Seurat FindClusters, 8 clusters were defined with a resolution of 0.3. FindMarkers was then run for each cluster, selecting gene markers (ident.1, only.pos=TRUE). UMAP density plots were obtained using the R package *Nebulosa* v1.8.0 and *scTransform* v0.4.1.

##### Expression correlation

To calculate the correlation of expression of two genes in a sample from single-cell-RNAseq data we employed *baredSC* v2.0.0 (Lopez-Delisle, et al, 2022). Here, the confidence interval of correlation is given as a percentage and the p-value, where *p* is the probability for the correlation coefficient to be negative, is the mean probability with the estimated standard deviation of this mean probability.

## Data availability

Sequencing data are available in the GEO repository under the accession number GSE259212.

## Acknowledgements

We thank Mylène Docquier, Brice Petit, Didier Chollet and Christelle Barraclough from the iGE3 sequencing facility. We thank Grégory Schneiter, Lan Tran and Cécile Gameiro from the Flow Cytrometry facility. We thank Olivier Fazio, Angélique Vincent and Fabrizio Thorel from the Transgenic facility. We thank Lucille Delisle for bioinformatic support. The computations were performed at University of Geneva using Baobab HPC service. This study was supported by grants from the Swiss National Science Foundation PP00P3_176802, PP00P3_210996, from the Novartis and Boninchi Foundations. M.K. lab is supported by the Canadian Institutes of Health Research grant CIHR 174989.

## Author contributions

G.A. conceived the project. O.B. and R.R.G. performed scRNA-seq preparations and analysis. O.B. and F.D. targeted and characterised the dCas9-P300 activator mESC clones and embryos. O.B. and A.R. performed mESC targetings, prepared the cells for tetraploid aggregation and performed WISH and skeletal preparations. O.B. performed embryo imaging, ChIP-seq, C-HiC and RNA-seq and analyses. M.K., F.G.-M. and C.G. provided the Eed knock out and control tissues for Capture-HiC. G.A. and O.B. wrote the manuscript with input from the remaining authors.

