## Supplementary Figures 1-7 for "Liebenberg syndrome severity arises from variations in *Pitx1* locus topology and ectopically transcribing cells"

### Supplementary information

#### Supplementary Figures

##### Supplementary Figure S1

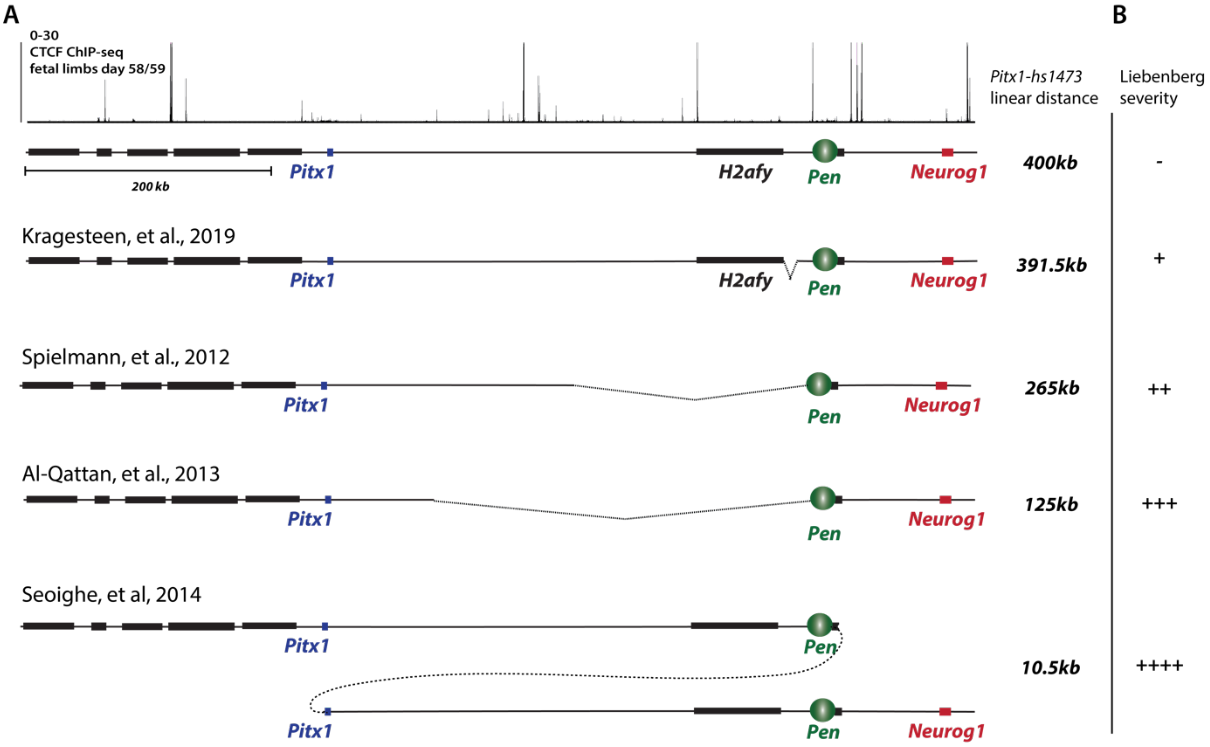


***Supplementary Figure S1:*** *We categorised the phenotypic description of Liebenberg anatomical features assigning a score based on the severity of the malformation: 0= not mentioned, 1= partially malformed and 2= severely malformed, taking also into account the physician's own comments on the severity of the cases (Supplementary Table 1). From this emerged that the more the distance between Pitx1 and Pen was reduced, the more severe was the manifestation of the condition (**Figure S2B**).*

**Supplementary Figure S2**

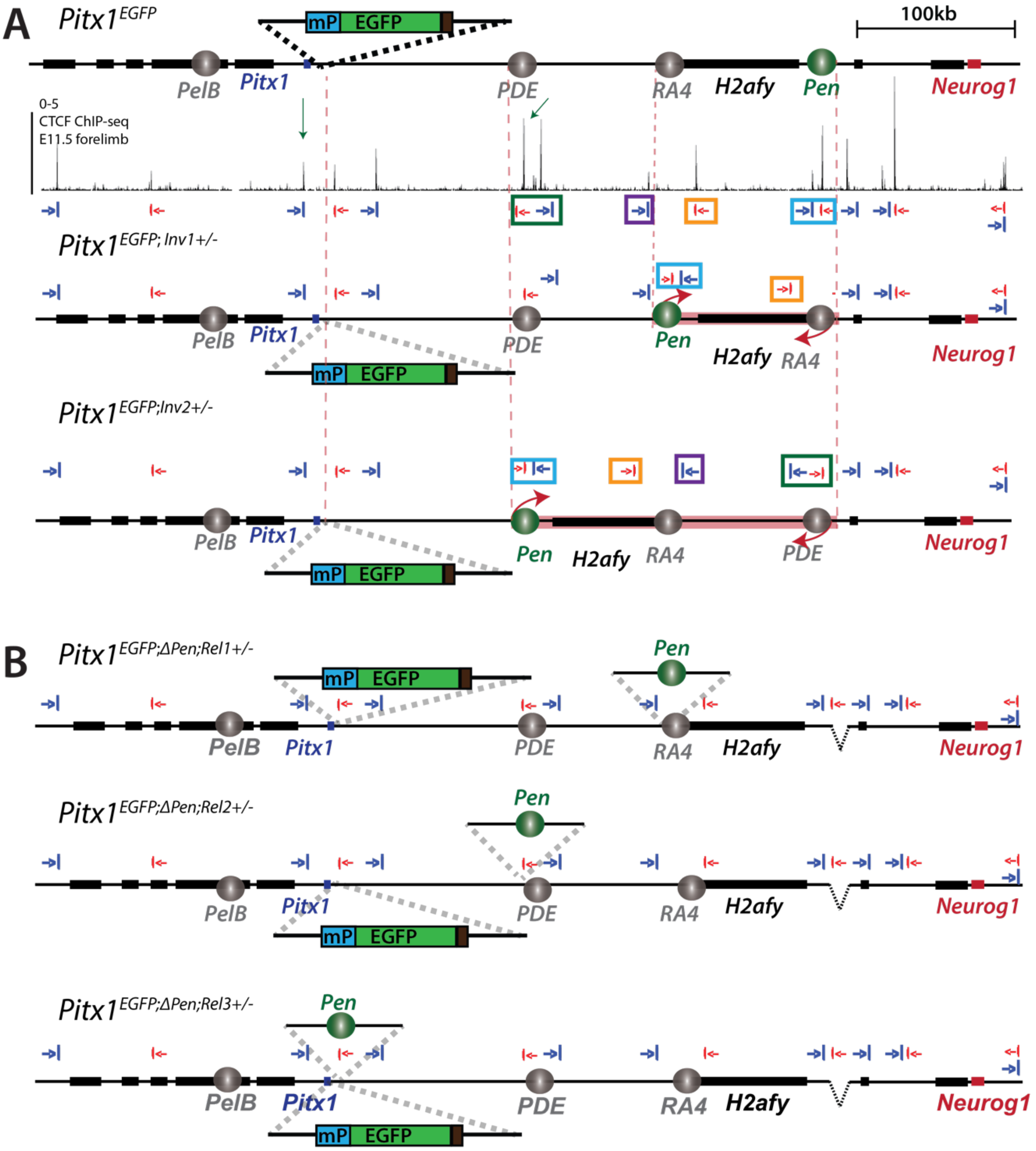


***Supplementary Figure S2: A.*** *ChIP-seq of CTCF binding at the Pitx1 locus shows several binding sites through the locus, in coloured boxes we highlight CTCF binding sites whose directionality is disrupted by the inversions Pitx1$^{EGFP;Inv1+/-}$ and Pitx1$^{EGFP;Inv2+/-}$, these CTCF sites are represented in the SVs at their new location and directionality. Red and blue arrows show CTCF binding site orientation, red dotted lines show the breakpoints of the SVs, green arrows on the CTCF ChIP-seq track indicates the two CTCF sites that form a stable loop between Pitx1 and PDE.* ***B.*** *A representation of the relocation of Pen approach (Pitx1$^{EGFP;\Delta Pen;Rel1+/-}$, Pitx1$^{EGFP;\Delta Pen;Rel2+/-}$ and Pitx1$^{EGFP;\Delta Pen;Rel3+/-}$) in a Pitx1$^{EGFP;\Delta Pen}$ background, where CTCF site orientation and position in the locus is not disrupted.*

#### Supplementary Figure S3

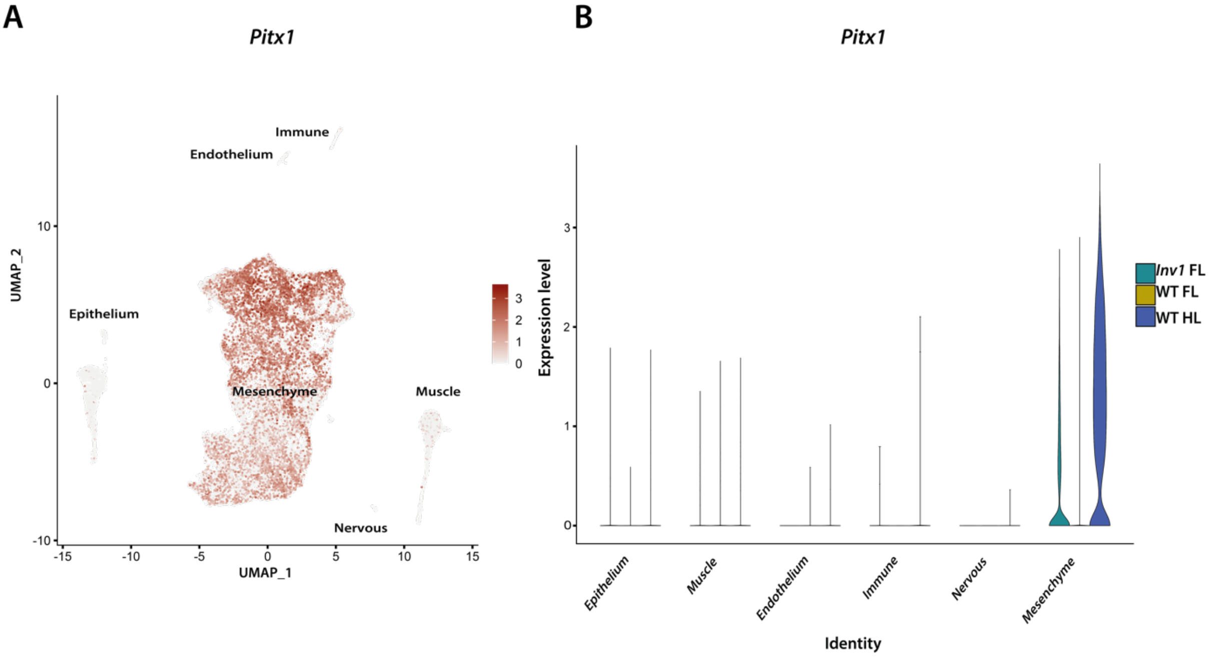


***Supplementary Figure S3:* A.** *UMAP representation of Pitx1 expression across cell types in wildtype E12.5 forelimbs, hindlimbs and E12.5 $Pitx1^{Inv1+/-}$ forelimbs.* **B.** *violin plots show expression levels of Pitx1 divided by limb identity and cell cluster, Pitx1 expression is restricted to mesenchymal cells and it is absent in wildtype forelimbs and present in wildtype hindlimbs* and *$Pitx1^{Inv1+/-}$ forelimbs (Inv1 FL).*

**Supplementary Figure S4**

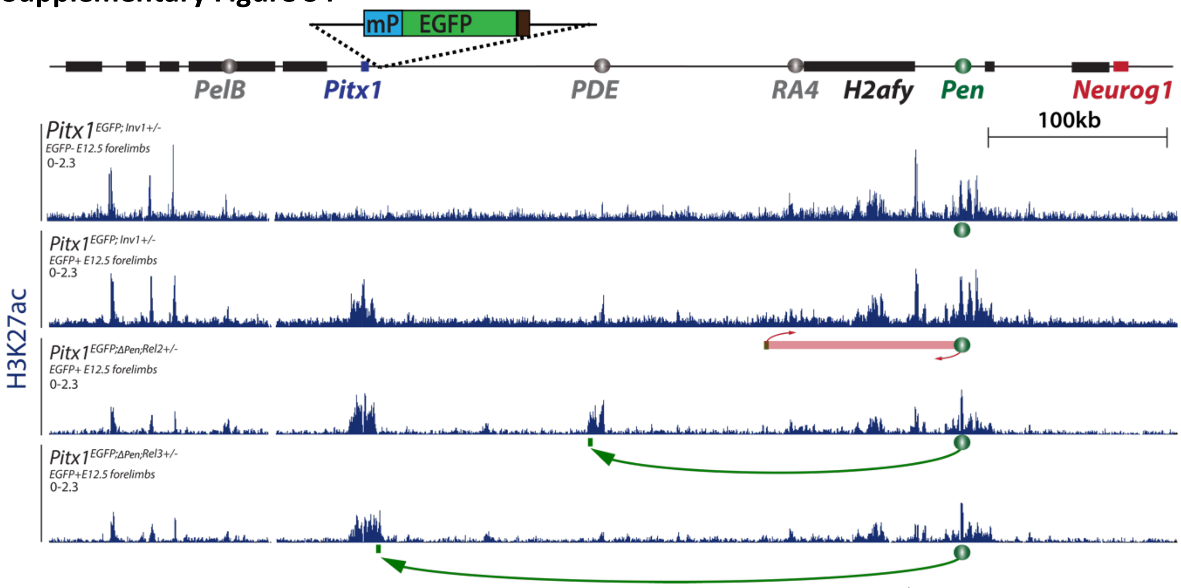


***Supplementary Figure S4:*** *A. H3K27ac ChIP-seq GFP- cells from Pitx1$^{EGFP;Inv1+/-}$, EGFP+ cells from, Pitx1$^{EGFP;Inv1+/-}$, Pitx1$^{EGFP;\Delta Pen;Rel2+/-}$ and Pitx1$^{EGFP;\Delta Pen;Rel3+/-}$ E12.5 forelimbs. Reads were mapped to a wildtype mm39 mouse reference genome. Note the real position of the Pen enhancers indicated by a green box within the inverted red box for the inversion, and by arrows for the two relocations.*

**Supplementary Figure S5**

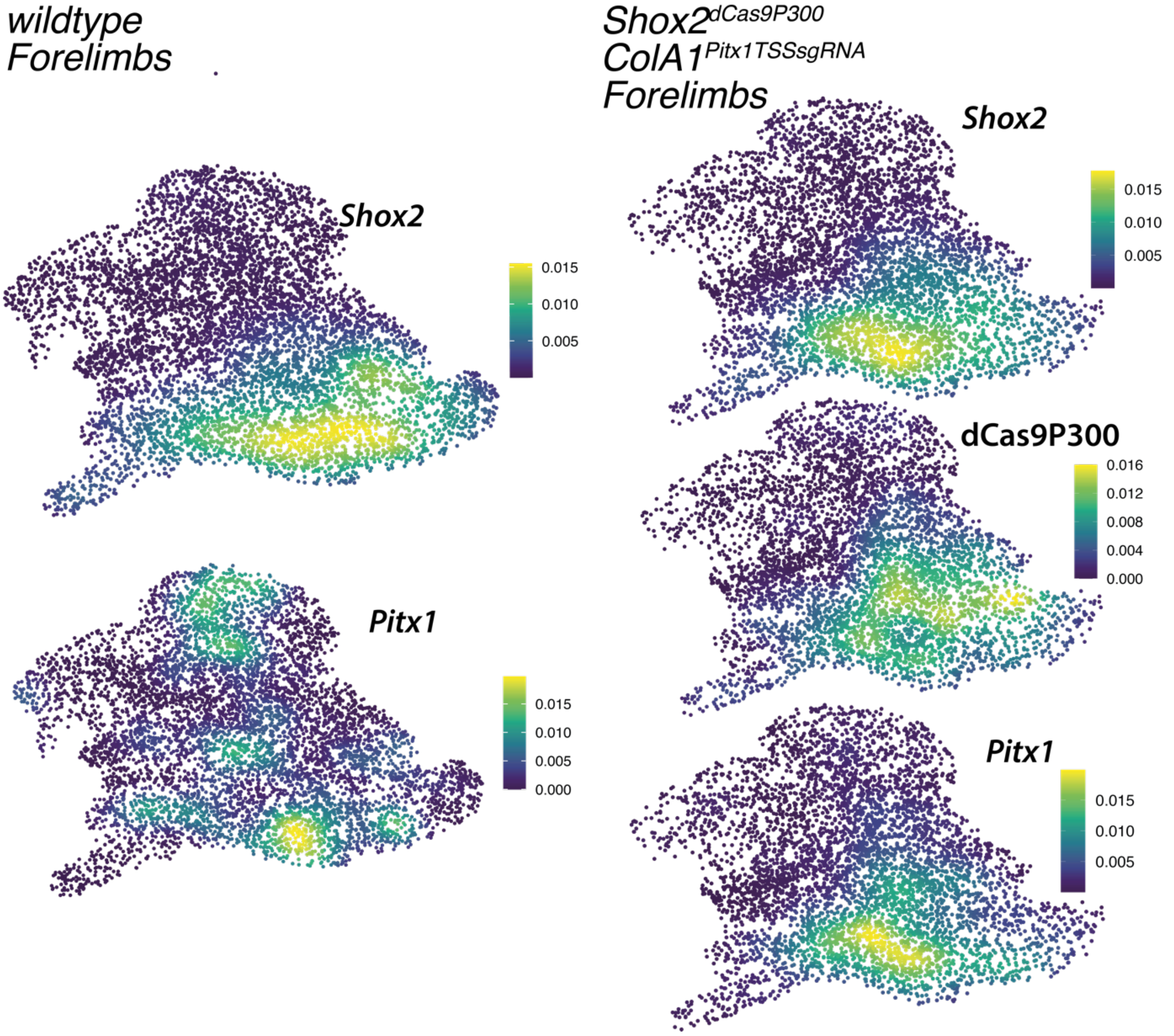


***Supplementary Figure S5****: Density plots of scRNA-seq data from wildtype forelimbs and $Shox2^{dCas9P300/+}$;$ColA1^{TSSsgR}$ forelimbs shows Shox2, Pitx1 and dCas9P300 expressing cells and co-localisation.*

#### Supplementary Figure S6

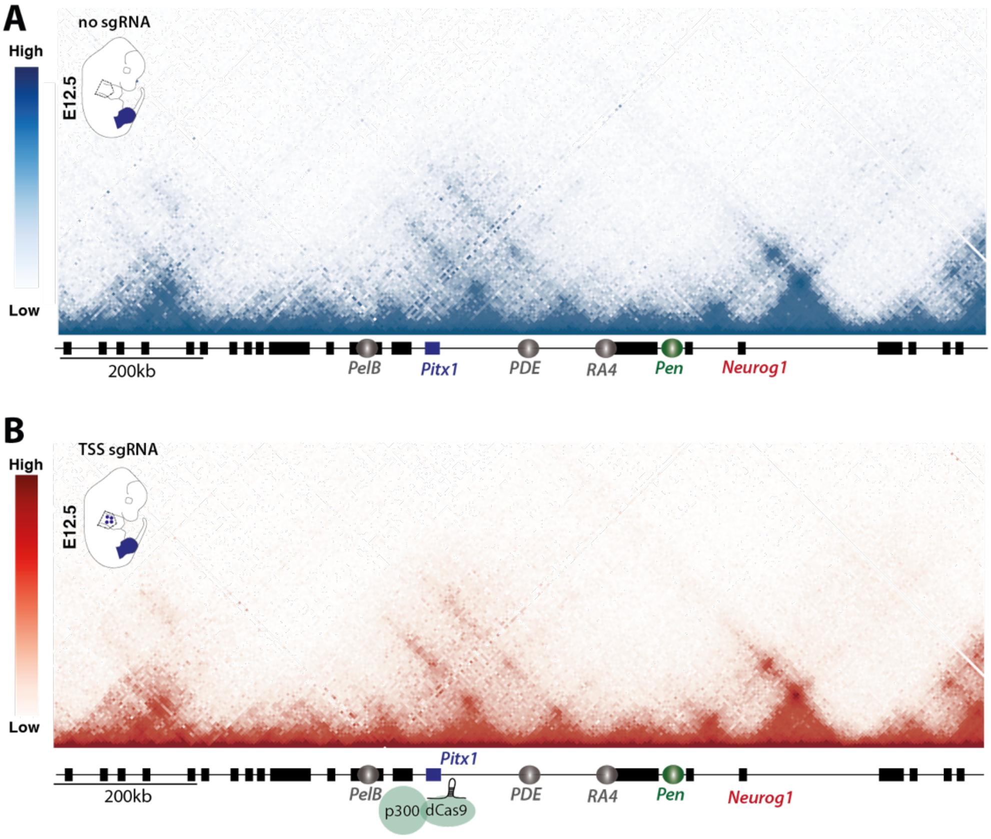


***Supplementary Figure S6:*** *C-HiC maps of (**A**) Shox2$^{dCas9P300/+}$;ColA1$^{+/+}$ (no sgRNA) and (**B**) Shox2$^{dCas9P300/+}$;ColA1$^{TSSsgR}$ (TSS sgRNA) E12.5 proximal forelimbs at the Pitx1 locus. Darker red or blue bins indicate stronger interaction frequencies as shown on the scale bars.*

*Supplementary Figure S7*

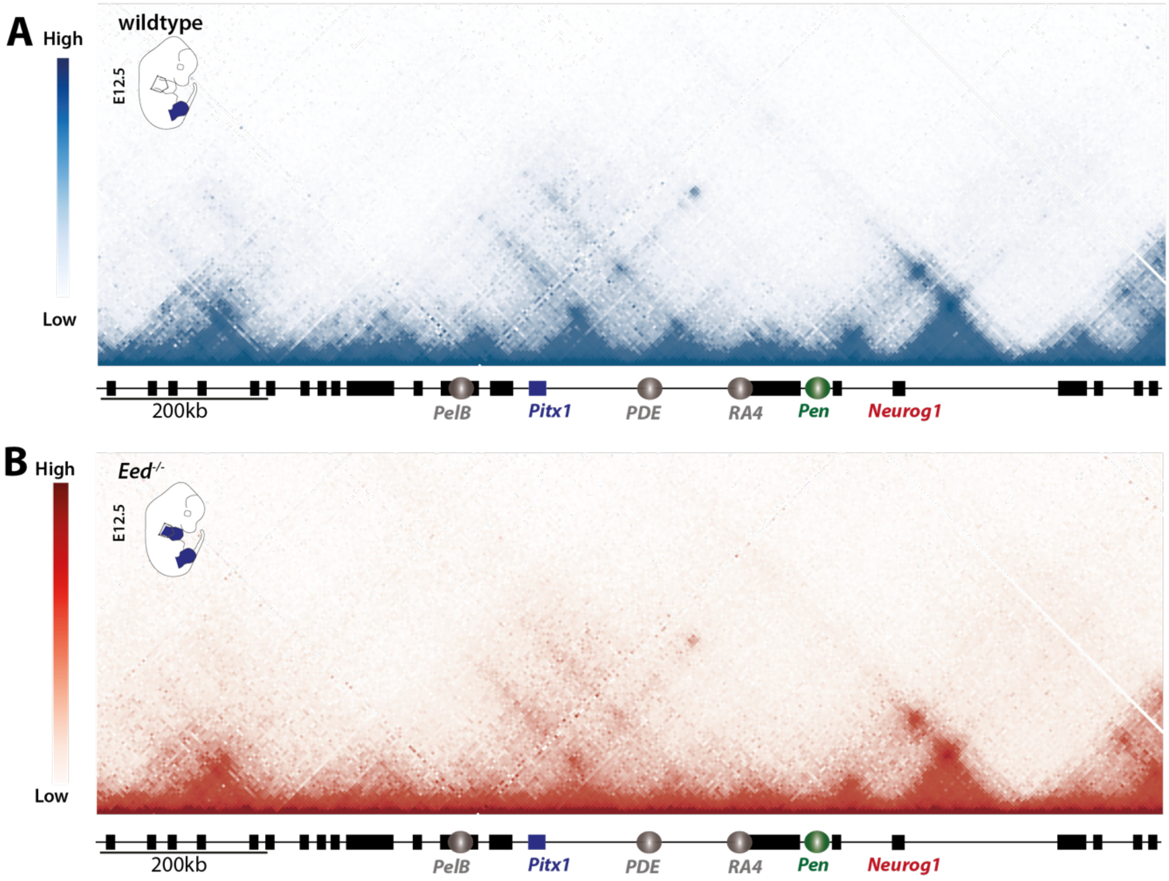


***Supplementary Figure S7:*** *C-HiC maps of (**A**) wildtype and (**B**) Prx1-Cre;Eed$^{flox/-}$ (Eed$^{-/-}$) E12.5 proximal forelimbs at the Pitx1 locus. Darker red and blue bins indicate stronger interaction frequencies as shown on the scale bars.*
